# Atmospheric nitrogen deposition and anthropogenic land use linked to changing fungal endophyte prevalence in cool-season grasses

**DOI:** 10.64898/2026.08.05.743054

**Authors:** Mallory N. Tucker, Tom E.X. Miller, Joshua C. Fowler

## Abstract

**Background and Aims:** Anthropogenic global change is altering the environmental stressors facing plants and their microbial symbionts. Changes in drought and temperature have received wide attention, but how other pervasive human impacts - land conversion for agricultural development and urbanization, and changes in nutrient conditions and pollutants - impact plant-microbe symbioses is relatively unknown. Here, we investigated how these anthropogenic global change drivers influence historic changes in the prevalence of widespread symbionts of grasses, *Epichloë* fungal endophytes.

**Methods:** We examined 8,739 seeds from 1,951 herbarium specimens collected between 1895 and 2019 for the presence of seed-transmitted *Epichloë* fungal endophytes in three grass host species (*Agrostis hyemalis, Agrostis perennans,* and *Elymus virginicus*). We hypothesized that the symbiosis provides fitness benefits under anthropogenic stresses (i.e. increased nitrogen deposition and land use change) that should translate to increased prevalence of the interaction among specimens exposed to those stresses.

**Key Results:** Anthropogenic stresses had contrasting effects on endophyte prevalence. Notably, among *Agrostis* host species, high nitrogen deposition was associated with high endophyte prevalence and with increasing trends in prevalence through time. We also found that highly urbanized landscapes were associated with reduced prevalence and negative temporal trends in endophyte prevalence across species. We also identified a weak positive relationship between agricultural land cover and average endophyte prevalence for *Elymus virginicus*, though temporal trends in prevalence did not differ between high and low levels of agricultural land cover.

**Conclusions:** Anthropogenic stressors influenced endophyte prevalence in diverse ways. While we found increasing prevalence in the face of nitrogen deposition, a sign of the potential resilience of the symbiosis, urban land cover was associated with declining endophyte prevalence, a sign that anthropogenic activity may contribute to a breakdown of the symbiosis.

## Introduction

Anthropogenic global change has driven broad shifts in the mean and variability of climate (Hansen and Stone 2016). On top of this shifting climate regime, many organisms occur in severely altered environments, including habitats converted for urban (Seto et al. 2013) and agricultural uses (Tilman et al. 2011), and in habitats affected by pollution and nutrient enrichment. At the same time as plants face these stressors, they engage with a variety of symbiotic fungal partnerships, including arbuscular mycorrhizae (Parniske 2008) and foliar endophytes (Rodriguez et al. 2009) that hold potential to ameliorate stress. How plants respond to these interactive global change drivers, and how symbioses influence host responses, are pressing questions.

Land use change, nitrogen deposition, and pollution are among the greatest direct threats to terrestrial plant diversity (Giam et al. 2010; Sage 2020). In addition to the direct loss of habitat and landscape connectivity, land use change alters the environments inhabited by plants and can reduce species’ resilience to changing climate (Foley et al. 2005; Oliver and Morecroft 2014). Urbanizing landscapes across the globe challenge plants with warmer temperatures and inconsistent water access due to impervious land cover (Johnson and Munshi-South 2017; Rivkin et al. 2019). Rising global temperatures combined with the heat island effect (the tendency for urban areas to have warmer temperatures compared to surrounding rural areas (Oke 1973)) may push species beyond their physiological thermal limits in urban environments (Kullberg et al. 2023). Alongside urbanization, agricultural intensification threatens wild plant populations (Newbold et al. 2015). Beyond habitat destruction, intensive agricultural practices have far-reaching indirect impacts including pollution with herbicides and pesticides (Rashid et al. 2010; Sun et al. 2018; Ilampooranan et al. 2022; Luna Juncal et al. 2023), as well as introducing increased pathogen loads in native plant communities (Susi and Laine 2021). Agricultural activities also contribute to eutrophication, the accumulation of nutrients such as nitrogen and phosphorus in soils and bodies of water (Payne et al. 2017). More broadly, agricultural practices, alongside fossil fuel combustion, contribute to atmospheric nitrogen deposition, which has more than doubled into terrestrial landscapes during the latter half of the 20th century (Galloway et al. 2004; Cleland and Harpole 2010; Nopmongcol et al. 2019). Nitrogen deposition rates vary greatly across regions, and have even declined in recent decades for some regions (Zhu et al. 2025). Despite this, nitrogen deposition rates remain above estimated pre-industrial levels (Engardt et al. 2017; Clark et al. 2018). While nitrogen deposition can benefit plant growth to a point, high deposition hinders plant growth through direct toxicity, and can indirectly alter biotic interactions within plant communities that compete over terrestrial nitrogen (Bobbink et al. 2010).

Benefits provided by fungal symbionts have the potential to mitigate the stress that confronts plant hosts under anthropogenically-driven global change (Rudgers et al. 2020; Schroeder et al. 2021). Many fungal symbionts have been shown to improve performance under drought (Gehring et al. 2017; Cheng et al. 2021) and warming (Staddon et al. 2004; Hector et al. 2022). These stress-tolerance benefits suggest that symbionts may enhance host resilience to exacerbated stresses in anthropogenic environments. The alternative expectation is that high stress increases the costs of maintaining the interaction, leading symbioses to break down, such as in the well-studied case of coral bleaching (Weis 2008). Forecasting the dynamics of plant species into the future requires mechanistic understanding of shifting costs and benefits of plant-fungal symbioses that support plant biodiversity (Fei et al. 2022).

Here we study how anthropogenic drivers influence plant symbioses with *Epichloë* foliar endophytes, ubiquitous fungal symbionts of grasses that are estimated to associate with up to 30% of cool-season grass species (Leuchtmann 1993). These endophytes are specialized, vertically-transmitted symbionts that live in the aboveground tissues of their hosts. While the interaction is obligate for the fungi, from the perspective of the host it is a facultative interaction that has been shown to reduce herbivory via the production of fungal alkaloids (Brem and Leuchtmann 2001; Rudgers et al. 2012; Gundel et al. 2020) and to enhance drought tolerance and nutrient uptake (Elmi and West 1995; Rudgers et al. 2009; Nagabhyru et al. 2013; Xu et al. 2017; Decunta et al. 2021). Because *Epichloë* fungi are primarily vertically-transmitted, symbiont effects that improve plant fitness should lead to increases in symbiont prevalence through time (Fine 1975; Gundel et al. 2008). The connection between host and symbiont fitness predicts that population-level endophyte prevalence should exist at some equilibrium that reflects the interaction’s fitness benefits given different environmental conditions, vertical transmission efficiency, and transient shifts in prevalence (Donald et al. 2021). Previous studies have documented considerable variation in mean endophyte prevalence across populations, and explained this variation as a result of symbiotic fitness benefits associated with broadscale environmental covariates (e.g. drought) (Afkhami 2012; Sneck et al. 2017; Li et al. 2026). What remains unclear is how other, non-climatic anthropogenic drivers influence the response of these interactions to global change.

*Epichloë* endophytes improve host responses to drought and temperature stress, yet there has been little research of the multiple stresses of land use change on grass-*Epichloë* symbioses (but see (König et al. 2018; Bastias et al. 2017)). Research in other systems provides examples of negative effects of land use change on plant-fungal symbioses. Surveys across anthropogenic habitat alteration and fragmentation found altered composition and function of soil microbial communities that potentially interact with plant hosts (Kiesewetter and Afkhami 2021), and an investigation of plant-mycorrhizal symbiosis along an urban land cover gradient showed disruption of colonization of tree species by their fungal symbionts (Tonn and Ibáñez 2017). Similarly, high nitrogen deposition also has the potential to disrupt plant-fungal symbioses. For example, for arbuscular mycorrhizal fungi that commonly improve nutrient uptake for hosts, nitrogen deposition is correlated with reduced root colonization, spore density, and hyphal length (Lilleskov et al. 2019). How nitrogen impacts *Epichloë* fungal symbioses has been less well studied, but the growth advantages of the interaction have been shown to decrease with the addition of nitrogen (Wang et al. 2018; Zhang et al. 2022), suggesting that improved nitrogen budgets are a benefit of foliar endophytes, not just root-associated symbioses. *Epichloë* fungi also produce nitrogen-rich secondary metabolites that underlie many of the benefits of the interaction. Nitrogen addition studies have documented contrasting effects on the concentration of these metabolites, sometimes leading to increased concentrations (Arechavaleta et al. 1992), and sometimes leading to reduced concentrations (Rasmussen et al. 2007). Thus, we expect nitrogen conditions to play an important role in mediating symbionts’ contributions to host responses to ongoing global change.

Herbaria, collections of plant specimens from across many years and wide geographic areas, provide unique insights into plants’ historic responses to changing environmental conditions (Lang et al. 2019). The large temporal and spatial extents provided by herbaria allow for investigations of plant responses to global change, including examinations of changes in herbivory (Meineke et al. 2019), phenology (Willis et al. 2017), and the spread of invasive species (Exposito-Alonso et al. 2018). For grass-*Epichloë* symbiosis, previous studies have used herbarium specimens to document host associations across the taxonomic diversity of grasses (White 1987). Fowler et al. (2025) used herbarium specimens to show that changes in climate drivers (seasonal precipitation and temperature) likely contributed to increasing endophyte prevalence through time, but did not entirely explain observed changes in endophyte prevalence, suggesting a role for non-climatic factors.

In this study, we investigated how anthropogenic land use and nitrogen deposition influence the prevalence of mutualistic *Epichloë* fungal endophytes. Using broadly-collected herbarium specimens (collected between 1895 and 2019) from three host species and data on land cover, nitrogen deposition, and climate drivers corresponding to specimen localities, we first asked how different anthropogenic drivers (urban land cover, agricultural land-use, and atmospheric nitrogen deposition) influence endophyte prevalence, averaging across collection years. Because endophytes can confer a fitness benefit to symbiotic hosts experiencing stress, we predicted that spatial variation in average endophyte prevalence would be positively associated with each anthropogenic driver after accounting for variation associated with climate. Then, we took advantage of the temporal nature of herbarium specimen data to explore whether trends in endophyte prevalence through time varied according to the intensity of each anthropogenic driver. We expected temporal shifts in endophyte prevalence to reflect responses to stresses which may be hidden in analyses that average across time when changes in anthropogenic stresses are rapid. Our study thus provides an assessment of the response of plant-fungal symbioses to diverse global change drivers, an important step in incorporating these biotic interactions into global change predictions.

## Materials and Methods

### Study Species

*Agrostis hyemalis, Agrostis perennans*, and *Elymus virginicus* are perennial cool-season grasses with broad native distributions across eastern North America. These species grow in partly shaded habitats in forest understories and savannas. *A. hyemalis*, in particular, can be commonly found in disturbed habitats such as roadsides. Each of these grass species host fungal endophytes in the genus *Epichloë* (*Epichloë amarillans* in *A. perennans* and *A. hyemalis* (Schardl and and Leuchtmann 1999) and *Epichloë elymi* in *E. virginicus* (White Jr. 1994)).

### Collection from herbaria

We collected seeds from specimens of the three focal host species (905 specimens of *A. hyemalis*, 366 specimens of *A. perennans*, and 680 specimens of *E. virginicus*) from nine herbaria within the United States (Table S1 provides a list of herbaria and specimen numbers). Our dataset included specimens spanning the Eastern continental United States collected as early as 1895 and as recently as 2019 (Fig. 1–2). While our herbarium surveys included specimens collected before 1895 (described in Fowler et al. 2025), we excluded pre-1895 specimens from this analysis because they were collected before the earliest year for which covariate data was available. To minimize tissue loss from specimens, we collected only small numbers of seeds (up to 10). *Epichloë* symbionts are predominantly vertically-transmitted through seeds from parent to offspring, however symbiotic plants can produce symbiont-free seeds via imperfect transmission (Afkhami and Rudgers 2008). Scoring multiple seeds per specimen as endophyte-symbiotic (E+) or symbiont-free (E-) accounts for this. The process of endophyte detection is described below.

**Fig. 1:**
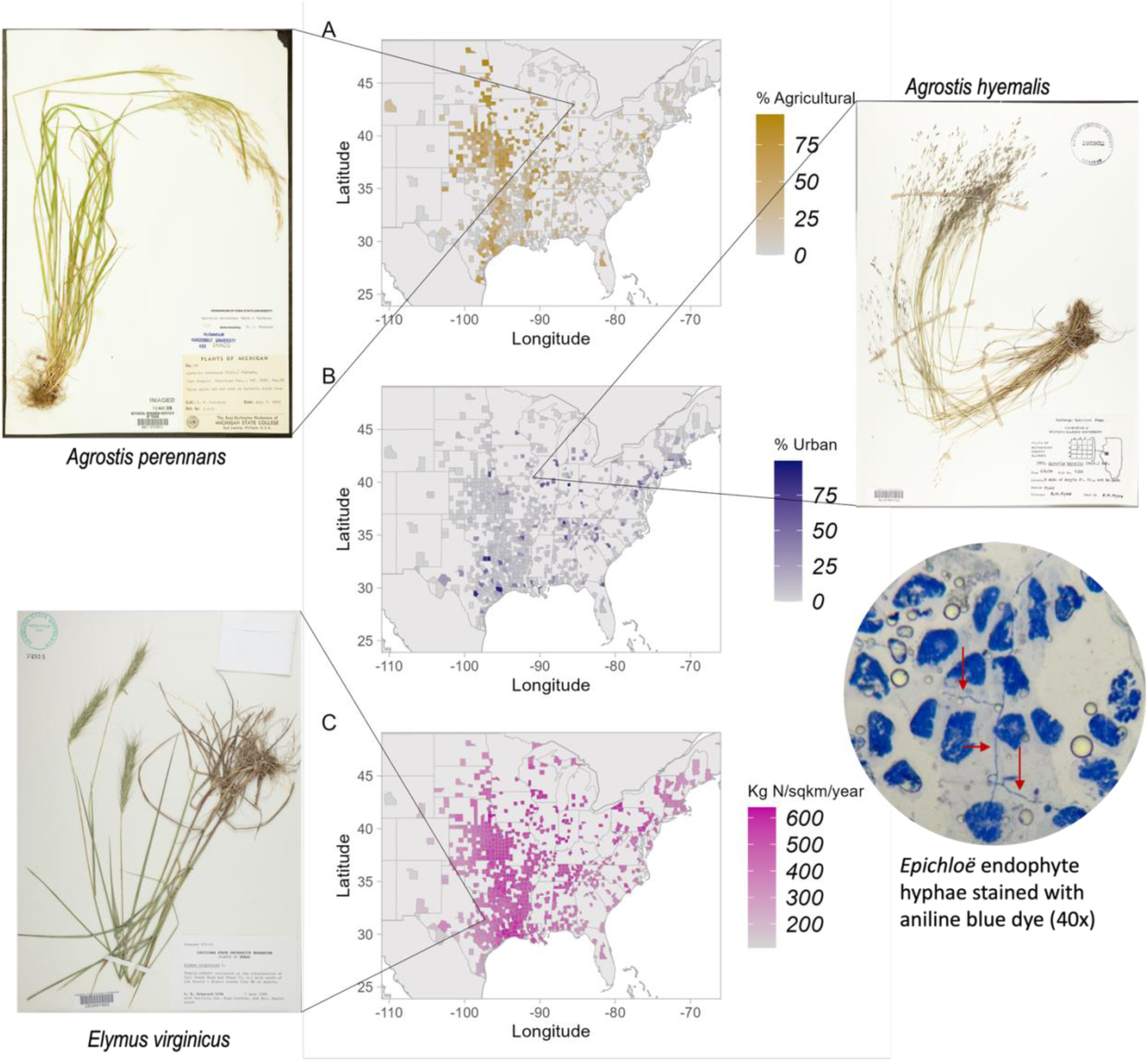
Geographic coverage of herbarium samples and anthropogenic global change drivers across the survey region. Shading depicts intensity of (A) percent agricultural land cover; (B) percent urban land cover; and (C) atmospheric nitrogen deposition (NO_3_ kg N/km^2^/month) within counties containing herbarium specimens surveyed for endophyte status. Photographs of *Agrostis perennans* and *Agrostis hyemalis* are taken from the Botanical Research Institute of Texas (CC0 1.0). Photograph of *Elymus virginicus* is taken from the Louisiana State University, Shirley C. Tucker Herbarium (CC BY-NC 3.0). Image of fungal endophyte hyphae modified from Brent Milton at Rice University.

**Fig. 2:**
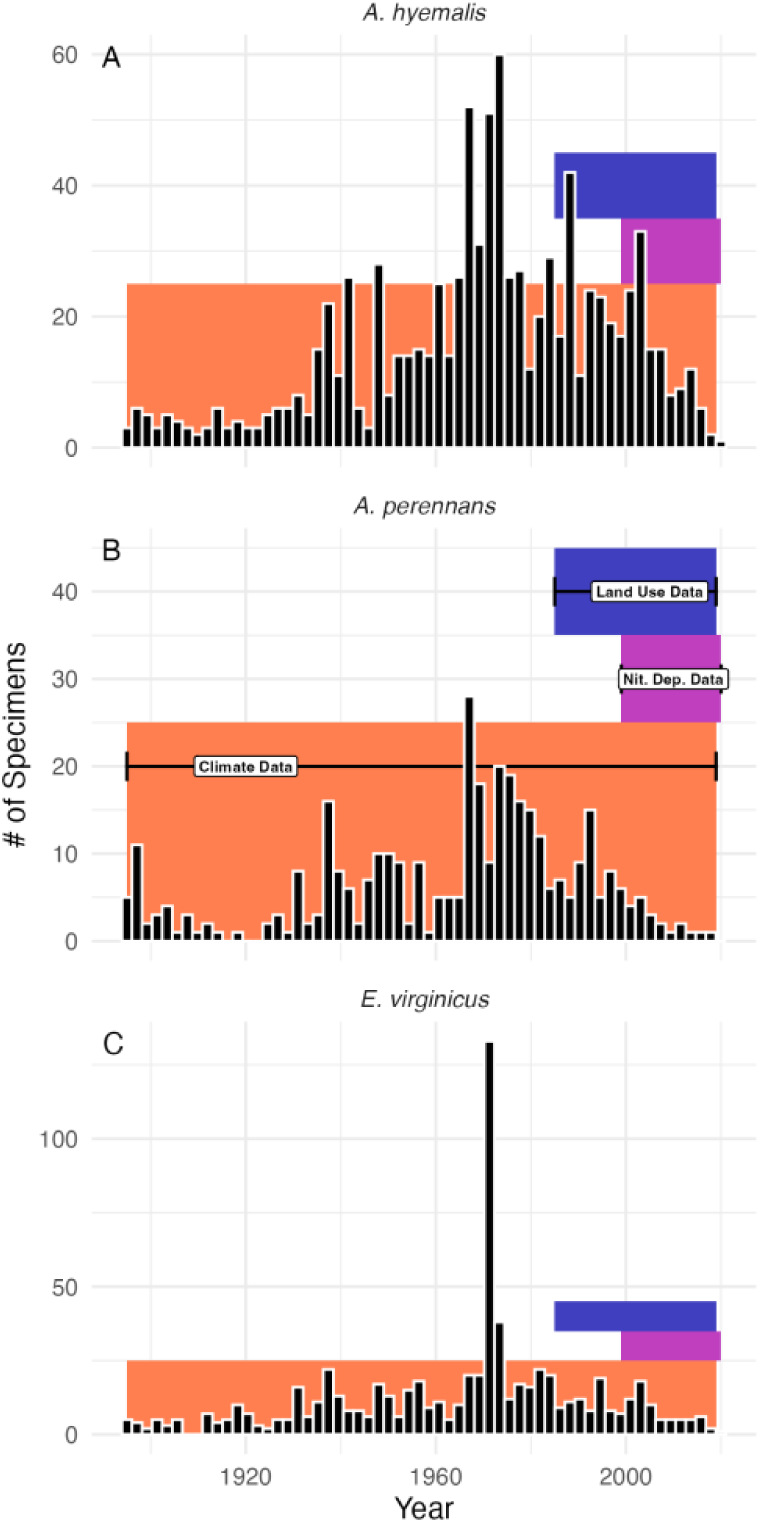
Temporal coverage of herbarium specimens and global change datasets across the survey region. Histograms show the count of herbarium specimens between 1895 and 2019 for (A) *Agrostis hyemalis* (B) *Agrostis perennans*, and (C) *Elymus virginicus*. Colored areas show the coverage of global change covariate datasets (orange: annual mean temperature and annual precipitation from the PRISM climate group (1895-2019); pink: Total Inorganic Nitrogen Deposition from USGS National Atmospheric Deposition Program (1999-2019); and purple: National Land Cover Database (1985-2019)).

To connect each specimen to its time and place of collection, we used digitized herbarium records from online databases, or for those not found in the databases, we transcribed metadata from herbarium labels. The locations of the specimens were georeferenced using the ggmap package (Kahle and Wickham 2013). Except where noted, all analyses were conducted in R version 4.3.3 (R Core Team 2024). We assigned coordinates to the nearest county or city centroid depending on available information. In most cases, specimens were assigned their county centroid (1491 specimens), and the rest were assigned their city centroid (460 specimens). The median georeferencing bounding box encompassed a large area (approximately 2200 km^2^). Geocoding may imply a highly precise location for some specimens (e.g., a less than 1 km^2^ area for some small municipalities). However, some specimens were collected in natural habitat nearby to, but not encompassed by, the bounding box around the georeferenced coordinates. We considered this range of georeferencing uncertainty in assigning covariate data to specimens, as described below, and explored how the choice of spatial scale impacted results.

### Endophyte detection

We stored seeds at -20 °C before they were examined microscopically. To determine the presence/absence of fungal endophytes, we first soaked seeds in a 10% NaOH solution to soften the seed coat. We dyed the seeds with lactic acid / aniline blue, squashed the seeds using a microscope slide cover, and then examined squashed seeds under a compound brightfield microscope at 100x-400x magnification for the presence of fungal endophytes (Bacon 2018). From the collected seeds that were successfully stained and scored (average number of scored seeds: 4.7 seeds per specimen for *A. hyemalis*, 4.2 seeds per specimen for *A. perennans*, and 3.8 seeds per specimen for *E. virginicus*), we recorded the number of seeds from each specimen in which we observed *Epichloë* hyphae (E+ for endophytes present, E- for endophytes absent). To capture uncertainty during the scoring process, we assigned endophyte status using both a “liberal” and a “conservative” score that indicate different thresholds of confidence in the endophyte assignment. Specifically, if we clearly observed at least one E+ seed, we would assign the specimen as E+ for both liberal and conservative scores. If we observed atypical endophyte morphology, or plausible endophytes obscured by poor staining, we assigned the plant a liberal score of E+ and a conservative score of E-. For those specimens where we confidently did not observe endophytes, we assigned matching E- liberal and conservative scores. In total, we collected the endophyte status of 1,951 grass specimens from 569 counties across the United States. Of these, 90.85% of *A. hyemalis,* 86.71% of *E. virginicus,* and 90.41% of *A. perennans* had matching liberal and conservative endophyte scores. Analyses presented below use the liberal scores, but we repeated all analyses using the conservative scores, which led to qualitatively similar results (Fig. S13-S14).

### Land Cover

We extracted land cover values from the National Land Cover Database (NLCD) (United States Geological Survey 2024) between 1985 and 2019 (Figs. 1, 2) with a 10 km radius buffer (314 km^2^) around the georeferenced coordinate points for each specimen’s collection coordinates using QGIS (QGIS Development Team 2024). To determine the adequacy of this buffer size, we repeated all subsequent analyses using 30 km radius buffers (2827 km^2^) which resulted in qualitatively similar model results (Fig. S7-S8). By summing pixels of different land cover types within the buffer for each year available to us in the landcover dataset, we calculated the percentage of urban land cover (Developed Open Space and Developed, Low, Medium, and High Intensity) and the percentage of agricultural land cover (Pasture/Hay and Cultivated Crops). We then calculated the average land cover for each category across years for each georeferenced collection location. The collection sites ranged from 0% agricultural to 95.60% agricultural and from 0.02% urban to 99.09% urban.

The statistical analyses presented below use spatially variable but temporally static land cover averages as predictor variables, although most herbarium specimens were collected before 1985 (Fig. 2), the first available year of land cover data. To assess the implications of using a single “snapshot” of land cover, we recreated our statistical analyses using year-specific land cover values with the subset of herbarium specimens for which year-specific values were available (n = 233 specimens). Across this subset of herbarium specimens, average land cover type (1985-2019) was tightly correlated with year-specific values at each collection location (Agricultural land cover: R^2^=0.989; Urban land cover: R^2^=0.992; Fig. S1). This correlation suggests greater variation in land cover across space than through time at a given location. It is likely however that the majority of land cover change occurred before 1985, and patterns of land use across space are not a direct measurement of land use change.

### Nitrogen Deposition

We used data on monthly inorganic nitrogen atmospheric wet deposition estimates for the conterminous United States (1999 through 2020) provided by the USGS National Atmospheric Deposition Program (Schmadel and Peterman 2023). After extracting the monthly total inorganic nitrogen deposition (kg N/km^2^/month) at the georeferenced coordinate points for each herbarium specimen within both a 10 km and 30 km buffer, we calculated the annual nitrogen deposition rate (kg N/km^2^/year) for each year between 1999-2020 (Fig. 2), as well as the average nitrogen deposition rate across the time period. Repeating all analyses with the 30 km buffer led to qualitatively similar results to the 10 km buffer (Figs. S16-S17). As above, we assessed the implications of a single temporal snapshot of nitrogen deposition by repeating our analysis using year-specific nitrogen deposition data with the subset of herbarium specimens for which year-specific values were available. Across this subset of specimens, year-specific values of nitrogen deposition were correlated with average nitrogen deposition (1999-2020) (R^2^=0.669; Fig. S1).

The collection sites ranged from 110.31 kg N/km^2^/year to 686.23 kg N/km^2^/year average nitrogen deposition. Across all locations, the mean nitrogen deposition was 472.94 kg N/km^2^/year. All analyses were conducted with total inorganic nitrogen, however we also repeated all analyses with extracted values from rasters of nitrate (NO_3_) and ammonium (NH4) ions, and found that it did not qualitatively impact results (Figs. S18 -S19).

### Climate covariates

While our central goal was to identify how endophyte prevalence relates to non-climatic anthropogenic global change drivers, we accounted for differences in climate conditions across the study region because regional differences in precipitation and seasonal warming have been linked to changing endophyte prevalence (Fowler et al. 2025). We extracted mean annual temperature and cumulative annual precipitation data from rasters provided by the PRISM climate group (PRISM, 2014). This dataset provides interpolated temperature and precipitation for each year between 1895 and 2019 (Fig. 2). We downloaded 800m resolution rasters and calculated mean values within 10 km and 30 km buffers around the georeferenced collection locations in each year of collection. Choice of buffer did not qualitatively alter results (Fig S16- S17).

### Statistical analysis

To test the role of anthropogenic global change drivers on endophyte prevalence, we constructed models including each of the focal drivers as predictor variables using an approximate Bayesian modeling framework, Integrated Nested Laplace Approximation, implemented in the INLA (Rue et al. 2009) and inlabru packages in R (Bachl et al. 2019). This framework is particularly useful because it allows computationally efficient implementations of spatially-structured random effects, accounting for the spatial non-independence of nearby herbarium specimens. Continuous spatially-structured random effects can be implemented using a stochastic partial differential equation (SPDE) approach (Lindgren et al. 2011; Bakka et al. 2018) where the covariance matrix of the spatial random effect is approximated using a Matérn covariance function. Each data point is assigned a location according to a mesh of non-overlapping triangles across the study area that defines the spatial decay between locations (Fig. S8). Further details of this approach applied to modeling endophyte symbiosis within herbarium specimens are provided in Fowler et al. (2025).

We modeled endophyte presence/absence as arising from a Bernoulli distribution, and all models shared the general form:

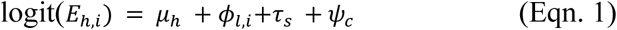

where the probability of endophyte positivity (*E*) of specimen (*i*) of host species (*h*) was modeled as a linear function of fixed effects (*μ*_ℎ_) comprising the effects of host identity and the focal global change variables (agricultural and urban land cover, nitrogen deposition, and climate covariates) along with random effects, including a spatially-structured random effect *φ* specific to each location (*l*) of each specimen, along with Gaussian distributed random effects (*τ* and *ψ*) accounting for the individual (*s*) who performed seed scoring and collector (*c*) who collected the specimen, respectively.

### Mean Prevalence Model

We first analyzed a model with spatially variable but temporally static anthropogenic driver variables to investigate their influence on average endophyte prevalence. This analysis relies on static, recent measurements of each anthropogenic driver (1980s-2020s) relative to the full temporal extent of the herbarium samples (1895-2019), reflecting data availability (Fig. 2). These covariates (land cover type and nitrogen deposition) are proxy variables that correspond to relatively more or less impacted habitats across the study region, but do not directly describe the rate of change over time in urbanization, agricultural development or nitrogen deposition. To account for the influence of climate on endophyte prevalence (as identified in Fowler et al. 2025), we also included climatic covariates (mean annual temperature and annual precipitation) from the year of collection.

In the Mean Prevalence Model, endophyte prevalence (the probability of endophyte positivity, *E*) of specimen (*i*) of host species (*h*) was modeled as described in Eqn. 1, with a fixed effects components *μ*_ℎ_:

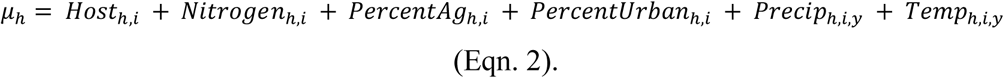

The model includes host species-specific intercepts and slopes for each global change driver of each observation (*i*). While nitrogen deposition and land cover covariates were average values, the mean annual temperature and precipitation are specific to the year of collection (*y*). In this Mean Prevalence Model, we are effectively asking how average endophyte prevalence across space is associated with spatial variation in these focal anthropogenic drivers while accounting for differences associated with climate, acknowledging that prevalence may not have reached equilibrium with rapid anthropogenic changes. We explore temporal dynamics of endophyte prevalence in the subsequent analysis, testing for interactions between temporal trends and anthropogenic drivers.

### Prevalence Trends Model

We next assessed whether trends in endophyte prevalence through time varied with the level of each global change driver. We predicted that prevalence should increase through time in response to stress and that stronger temporal trends should also be a signal of global change responses that might be obscured in the analysis of static anthropogenic drivers. Therefore, we modeled endophyte prevalence in a manner similar to Eqn. 1, however we included parameters describing host-species specific temporal trends and their interaction with our focal global change drivers. Here, the fixed effects component of the model, *μ*_ℎ_, is given by:

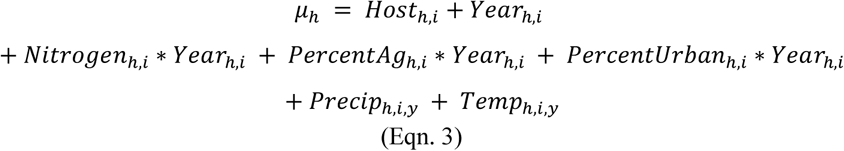

where the interaction terms between collection year and the global change drivers allowed us to evaluate temporal trends in prevalence across levels of each global change driver. All other parameters were as described in Eqn. 1.

To evaluate the effect of each anthropogenic driver on endophyte prevalence, we visualized conditional predictions across the range of each covariate while holding others at their mean value and also evaluated posterior probabilities of effect for the main and interactive effects. We evaluated model fit with graphical posterior predictive checks (Fig. S19).

## Results

### Anthropogenic drivers of endophyte prevalence

Overall, mean endophyte prevalence was 62.1% in *A. hyemalis* (Mean = 62.1%, SD = 23.6%), 63.4% in A. perennans (M = 63.4, SD = 23.3%), and 57.7% in *E. virginicus* (M = 57.7, SD = 24.4%) within herbarium specimens. Analysis of the Mean Prevalence Model revealed support for anthropogenic drivers of mean endophyte prevalence. We found mixed associations between agricultural land cover and endophyte prevalence across host species (Fig. 3 a-c). In particular, there was greater than 99% posterior probability that high levels of agricultural land cover were associated with high endophyte prevalence in *E. virginicus* (*posterior median* = 0.009, *95% CI* = [0.002, 0.017]; Fig. S10). This effect equated to a 43% increase in endophyte prevalence (from 47% to 67% prevalence) across the full gradient of agricultural land cover. For *A. hyemalis* and *A. perennans*, there was little evidence of an effect of agriculture with a 63% probability of a negative slope (*median* = -0.001, *CI* = [-0.007,0.006]) and 64% probability of a positive slope (*median* = -0.002, *CI* = [-0.009,0.013]) respectively (Fig. S10).

**Fig 3:**
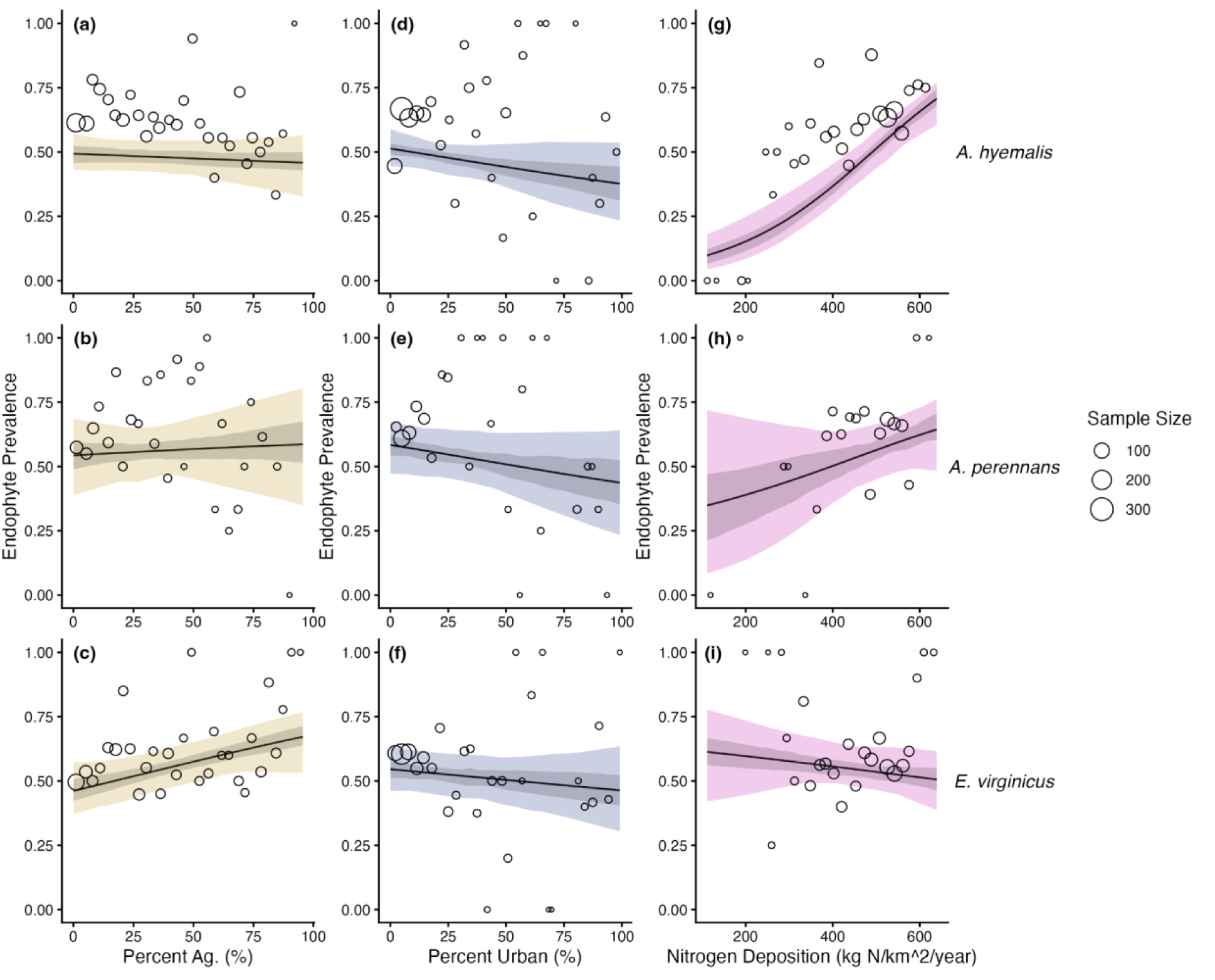
Relationships between anthropogenic global change drivers and *Epichloë* endophyte prevalence in three host species (rows), *A. hyemalis*, *A. perennans*, and *E. virginicus*. Lines depict predicted mean relationship between covariate and endophyte prevalence along with shaded 50% and 95% credible intervals for percent agricultural land cover (panels a-c; yellow), percent urban land cover (panels d-f; blue), and total inorganic atmospheric nitrogen deposition (panels g-i; pink). Points represent average endophyte prevalence as binned means across each covariate for each species with point size proportional to bin size.

Evaluating relationships between urban land cover and endophyte prevalence, we found weak, slightly negative effects consistently across host species (Fig. 3 d-f). For *A. hyemalis,* there was a 88% posterior probability of a negative relationship (*median* = -0.005, *CI* = [-0.013,0.003]; Fig. S10), with average endophyte prevalence shifting from 51% to 40% across levels of urban land cover, a 22% decrease. Similarly, there was a 82% probability of a negative relationship for *A. perennans* (*median* = -0.006, *CI* = [-0.018,0.006]; Fig. S10), and a 76% probability of a negative relationship for *E. virginicus* (*median* = -0.003, *CI* = [-0.012,0.005]; Fig. S10), effects which were of similar magnitude (16% and 26% decrease in endophyte prevalence respectively).

Additionally, our analysis indicated moderate to strong relationships between nitrogen deposition and endophyte prevalence across host species (Fig. 3 g-i). In *A. hyemalis*, predicted mean endophyte prevalence increased from 10% to 71% across the nitrogen deposition rates represented in our data, with greater than 99% posterior probability of a positive slope (*median* = 0.006, *CI* = [0.004,0.009]; Fig. S11). For *A. perennans*, the positive relationship was slightly weaker and more uncertain, increasing from 36% to 64% prevalence with an 86% probability of a positive slope (*median* = 0.002, *CI* = [-0.001,0.006]; Fig S11). In *E. virginicus*, we found a weak negative relationship corresponding to a shift in endophyte positivity from 60% to 51% prevalence with a 74% posterior probability of a negative slope) (*median* = -0.001, *CI* = [- 0.003,0.002]; Fig S10).

Notably, these conditional predictions reveal variation in endophyte prevalence explained by anthropogenic drivers after accounting for climate. We estimated a positive effect of precipitation for each species (ranging from 64% to 81% posterior probability of positive relationships)(*A. hyemalis*: (*median* = 0.0002, *CI* = [-0.0004,0.0007]), *A. perennans*: (*median* = .0002, *CI* = [-0.0008,0.0012]), *E. virginicus*: (*median* = 0.0003, *CI* = [-0.0003,0.0009]); Fig. S10), and both positive and negative effects of temperature on endophyte prevalence (86% probability positive for *A. hyemalis* (*median* = 0.030, *CI* = [-0.027,0.081]) and 98% probability negative for *E. virginicus* (*median* = -0.054, *CI* = [-0.103,-0.001]), Fig. S10). Correlations between focal anthropogenic drivers and climate covariates were generally low (Fig S5-S6), with the highest correlation coefficient value of 0.33 for the relationship between annual precipitation and nitrogen deposition (Fig. S5). Variance inflation factors were below 1.5 for all covariates (Fig. S7) suggesting no problematic collinearity.

### Influence of anthropogenic drivers on temporal trends in endophyte prevalence

Over the past century, endophyte prevalence increased on average for each of the three host species. After accounting for changes over this period associated with changes in climate drivers, endophyte prevalence increased by 17.5 percentage points per century (p.p./century) for *A. hyemalis* (*median* = 17.5%, *CI* = [-8.1%,40.3%]), 3.6 p.p./century for *A. perennans* (*median* = 3.6%, *CI* = [-35.0%,42.8%]), and 2.4 p.p./century for *E. virginicus* (*median* = 2.4%, *CI* = [- 27.7%,34.0%]).

Anthropogenic stressors contributed to substantial variation in temporal trends. We found generally increasing trends regardless of level of agricultural land cover (probability of positive trends: greater than 87%, 66%, and 69% for *A. hyemalis*, *A. perennans*, and *E. virginicus* respectively (Fig. 4). For example, for *A. hyemalis* specimens in low agricultural land cover, endophyte prevalence increased by 19.8 p.p./century while in highly agricultural habitats, endophyte prevalence increased by 24.8 p.p./century, with overlapping 95% credible intervals (low *CI* = [-5.0%,40.4%]; high *CI* = [-14.7%,55.0%]; Fig. 4a).

**Fig. 4:**
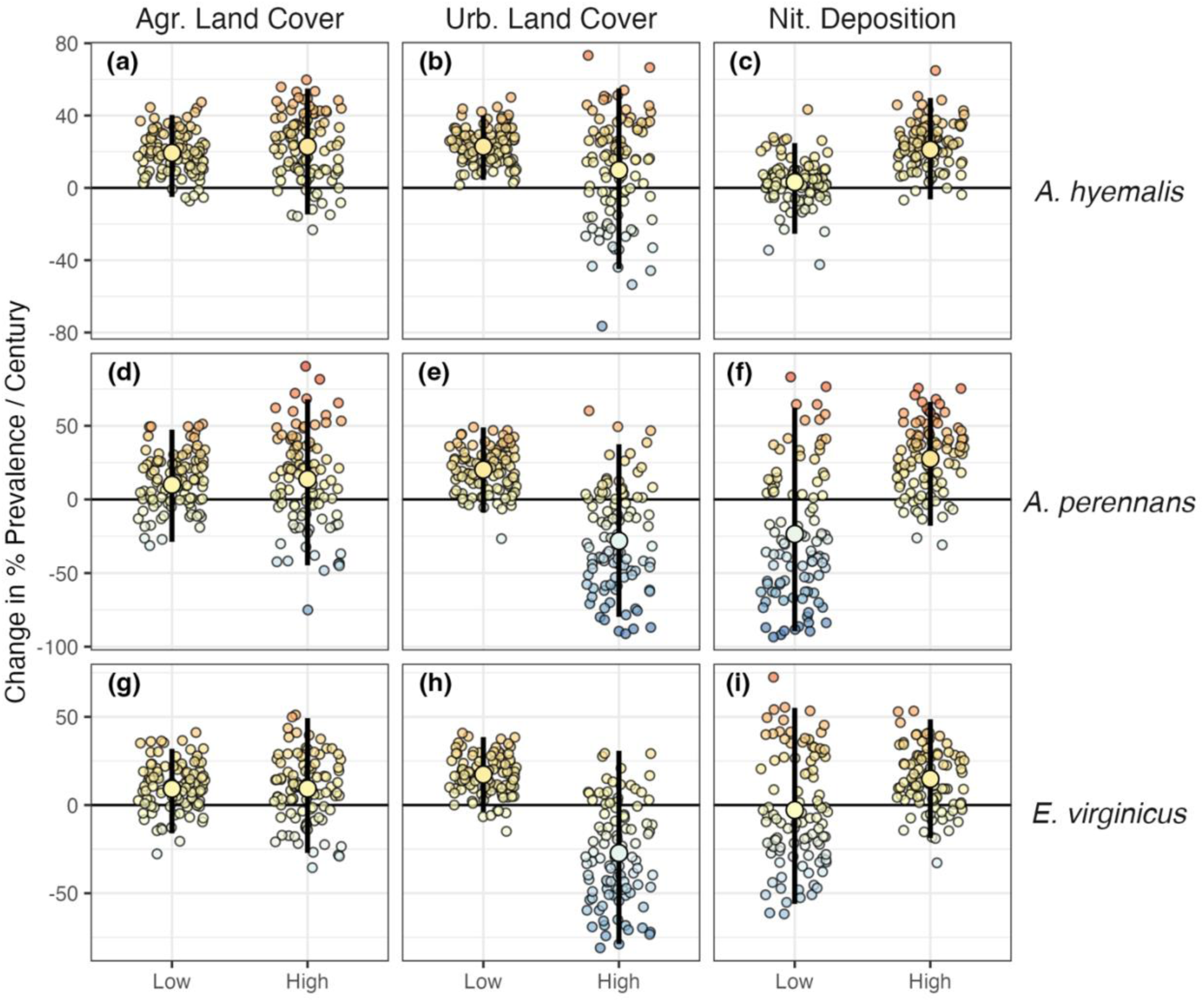
Anthropogenic drivers mediate temporal trends in endophyte prevalence. Panels show conditional predictions of change in endophyte prevalence per century across the minimum and maximum values of each anthropogenic driver (agricultural land use (a, d, g), urban land use (b, e, h), and nitrogen deposition (c, f, i) for each species (*Agrostis hyemalis*, *Agrostis perennans*, and *Elymus virginicus*). Central points and lines represent posterior means along with 95% credible intervals. Small point clouds are 100 posterior samples denoting uncertainty. Shading in points depicts the mean rate of change per century in endophyte prevalence.

Highly urban habitats were associated with declining endophyte prevalence for *A. perennans* and *E. virginicus*. Trends in endophyte prevalence shifted from 20.7 p.p. increases per century (90% probability of positive trend; *median* = 20.7%, *CI* = [-8.9%,48.9%]; Fig 3e) under low urban land use to 29.9 p.p. declines per century (82% probability of negative trend; *median* = -29.9%, *CI* = [-79.8%,37.5%]; Fig. 4e) under high urban land use, with a shift of similar magnitude for *E. virginicus* (Fig. 4h). Highly urban conditions also dampened positive trends in endophyte prevalence for *A. hyemalis*; strongly increasing prevalence under low urban land cover (99% probability of positive trend; *median* = 22.7%, *CI* = [4.4%,40.1%]; Fig 3b) weakened under high urban land cover (66% probability of positive trend; *median* = 12.1%, *CI* = [-44.9%,55.1%]; Fig 3b).

Finally, high nitrogen deposition was associated with increasing trends in endophyte prevalence, particularly for the *Agrostis* species. Under high nitrogen deposition, prevalence increased 21.4 p.p./century for *A. hyemalis* (94% probability of positive trend; *median* = 21.1%, *CI* = [-6.3%,49.7%]; Fig 3c) and 28.6 p.p./century for *A. perennans* (86% probability of positive trend; *median* = 28.6%, *CI* = [-17.9%,66.3%]; Fig 3f). These trends reflect substantial changes in endophyte prevalence over time. For example, the model predicted a rise in endophyte prevalence under high nitrogen deposition from 43% to 76% prevalence among specimens of *A. perennans* between the years 1900 and 2000.

A visualization of the rate of change in endophyte prevalence across the full covariate space is provided in the supplemental material (Fig. S11-S12). We note that these temporal trends reflect change in prevalence associated with each anthropogenic driver after accounting for precipitation and temperature.

While the above results come from analysis of static “snapshot” covariate data, they compare favorably to analysis of year-specific covariate data. In general, we found that the posterior estimates concur between snapshot and year-specific data for both the Mean Prevalence and the Prevalence Trends Models (Supplemental Methods, Figs. S2-S4). This supplemental analysis uses a much smaller subset of the data (n = 244 specimens), and as a result, parameter credible intervals are wider and overlap zero. Despite this, these supplemental analyses generally agree with the direction of effects found in the main analyses. For example, with year-specific covariates, we estimated weak positive effects of nitrogen deposition on average prevalence (Fig. S2) and on temporal trends in prevalence (Fig. S3) for *A. hyemalis*. Some parameter estimates differed, particularly for the Temporal Trends model. For example, while we found a negative effect of urban land cover on temporal trends for *E. virginicus* in our central analysis (Fig. 4h), analysis of year specific data resulted in weakly positive posterior trends (Fig. S3). Comparisons for *A. perennans* are limited by small sample size (n = 22 specimens).

## Discussion

Our analysis of herbarium specimens from across the eastern United States revealed strongly supported relationships between plant-fungal symbiosis and anthropogenic drivers of global change related to land use and eutrophication. Because these symbionts are predominantly vertically transmitted within host populations, we expect that the average prevalence of the interaction and changes in endophyte prevalence are indicators of the fitness consequences of the symbiosis and the response of the symbiosis to global change drivers. A major takeaway is that nitrogen deposition appears to be associated with high and increasing prevalence of this symbiotic mutualism. Yet, we also found evidence that the prevalence of the symbiosis may be declining under the combined stresses of anthropogenic activity, particularly in urban environments.

Atmospheric nitrogen deposition, in particular, appears to strengthen the symbiosis between grass hosts and their vertically-transmitted *Epichloë* endophytes. Regions with high levels of nitrogen deposition were associated with high average endophyte prevalence, particularly for the *Agrostis* species. High nitrogen deposition was also associated with increasing endophyte prevalence over time. Notably, this result held when controlling for variation in precipitation, which is correlated with nitrogen deposition (R^2^ = 0.66) and itself contributes to increasing endophyte prevalence in these host species (Fig S12). These results suggest that *Epichloë* symbiosis improves host fitness relative to nonsymbiotic hosts under high nitrogen deposition. This fitness benefit could arise either when symbionts provide resilience to stress from excess nitrogen deposition, or when symbiotic plants are better able to take advantage of added nitrogen in nitrogen limited environments.

The ability to uptake nitrogen and nitrogen use efficiency are important components of plants’ physiological responses to increased levels of atmospheric nitrogen deposition (Gastal and Lemaire 2002; Lemaire 2021). Fungal endophytes, by utilizing nitrogen and modifying host plant physiology, may play an important role in host responses to nitrogen stress. Limited research has been conducted on the interplay between the physiology of nitrogen uptake and the role of *Epichloë* endophytes, but we suspect that this could explain varying responses to increased nitrogen deposition between grass hosts. Inter- and intra-specific variation in root nitrate uptake rate, leaf N content, and nitrogen use efficiency in response to soil NO_3_ been observed in related *Agrostis* species (DaCosta et al. 2022), indicating that species differences in response to increased soil nitrogen could determine the ecological effects of anthropogenic atmospheric nitrogen deposition. Moreover, the quantity and production of nitrogen-rich alkaloids differs across endophyte strains (Saikkonen et al. 2013), and there is evidence that some *Epichloë* endophytes may confer benefits by directly interacting with nitrogen-associated enzyme pathways (Helander et al. 2016; Wang et al. 2018). Thus, we suspect that the combination of host- and endophyte- specific traits may produce different responses in host species to nitrogen addition. Alongside the responses directly elicited by nitrogen, nitrogen deposition can alter herbivore population densities (Graff et al. 2020), plant-pathogen interactions (Easterday et al. 2022), and plant community composition (Henning et al. 2021), each of which also have the potential to increase the importance of grass-endophyte symbiosis, mediated by well-known anti-herbivore compounds produced by *Epichloë* (Crawford et al. 2010) and through endophyte-enhanced competitive abilities (Vázquez-de-Aldana et al. 2013).

Agricultural land use was positively associated with higher average endophyte prevalence only for *E. virginicus*, but we did not find clear evidence that greater agricultural land use favors higher prevalence for the *Agrostis* species. Similarly, agricultural land use was not associated with changes in temporal trends in prevalence. The primary impact of agricultural land use is habitat loss, which is likely to similarly impact symbiotic and non-symbiotic plants. However, we expected that *Epichloë* symbiosis has the potential to mitigate some stresses associated with agricultural land use: increased disturbance and edge effects, changes in soil salinity, and exposure to pollutants, as well as natural enemies that disperse across habitat edges between agricultural and intact land (Rand et al. 2006; Inclán et al. 2016; Mazón et al. 2024). These agriculture-associated stresses may differ substantially across different agricultural habitats in ways that are not captured in remotely-sensed land use categories. The broad spatial scale of our analysis – using georeferenced herbarium specimens and agricultural land cover as a proxy for these various stresses and biotic interactions relevant to individuals at local scales – potentially limits our ability to detect effects on endophyte prevalence. In addition, while *Epichloë* fungi have been shown to protect hosts against herbivores, pests, and pathogens (Porter et al. 1981; Shiba et al. 2015; Schardl, Young, et al. 2013; Kou et al. 2021), the benefits of *Epichloë* symbiosis often depend on species and genotype-level differences of each interacting partner. Herbivore protection mediated by *Epichloë* symbionts has been evaluated previously in these three host species; only the endophyte of *A. perennans* has been shown to reduce insect herbivore damage and impair herbivore performance through the production of toxic alkaloids (Crawford et al. 2010). However these herbivory assays examined plants reared from singular host populations. Because population-specific differences in alkaloid concentrations and in the genetic loci associated with the production of different alkaloid types are common among *Epichloë* symbioses (Schardl et al. 2013; Sneck et al. 2019), it would be particularly interesting to pair studies of changing symbiont prevalence with genotyping for the genes that underlie herbivory defense.

Finally, our results suggest that urban land-use change may be a potential driver of deterioration in these plant-fungal symbiosis. High levels of urban land cover were only moderately associated with reduced endophyte prevalence on average. However this effect was consistent across species, and we also found that highly urban habitats were likely to be associated with declining endophyte prevalence over time for both *E. virginicus* and *A. perennans*. Urban environments induce unique stresses for plants - increased temperature and drought (Knapp 2010; Huang et al. 2024), along with disruptions to dispersal and reproduction (Otto 2018; Miles et al. 2019). Reductions in fitness associated with urban land use may align with previous research that finds that the benefits of partnership with fungal endophytes, for *E. virginicus* in particular, are reduced under drought stress (Rudgers and Swafford 2009). While this interpretation conflicts with the prevailing belief that *Epichloë* can provide drought tolerance benefits to their hosts, drought tolerance benefits, like herbivore defense, can depend upon species and even genotype-specific differences (Decunta et al. 2021). Moreover, it is possible that urban environments are especially stressful for the host species, combining drought and temperature stress along with pollutants such as ozone (Paoletti et al. 2014), that potentially weaken the hosts’ ability to partner with symbionts (Bastías and Gundel 2023). This reiterates the need for experiments to disentangle mechanistically the response of plant-fungal symbioses to multiple interacting global change drivers (Rillig et al. 2019), following up on the associations we identified in this study.

Our use of herbarium specimens also highlights key challenges in global change research more generally: appropriately connecting and interpreting the temporal and spatial scales of different sources of global change data, including data that provide information on human influences beyond commonly available gridded climatic covariates (Frans and Liu 2024). Recent rapid changes in the environment, such as the dramatic changes in nitrogen deposition in the last 50 years and the expansion of urban land-use, likely put host-microbe symbiota distributions in disequilibrium with their environmental optima. While ecological niche models estimate environment-occurrence relationships by averaging over spatial occurrences across many years (Araújo et al. 2005; Gallien et al. 2012), some responses may lag behind observed changes in global change drivers. Identifying shifts in response to rapidly changing environmental drivers requires analyses that are spatially accurate and temporally-explicit or that incorporate ecological dynamics (Franklin 2010; Merow et al. 2011; Carlson et al. 2022). A response lag may be one reason we found stronger evidence of a relationship between trends in endophyte prevalence and nitrogen deposition than between nitrogen deposition and average endophyte prevalence.

At the same time, due to data availability, most of the analyses in this study were performed with static estimates of anthropogenic global change drivers taken from a relatively recent period relative to the extent of the specimen dataset. To make the causal interpretation that change in a given driver is driving change in endophyte prevalence would require making an important assumption: that regions with high values of a given driver (e.g. urban land cover) are those that have experienced the greatest change in that driver. We expect that while this may be true in some cases, such as for nitrogen deposition which has rapidly increased more than two-fold on average over the last 50 years, it is likely not always so. Have regions that were rural or those that were urban in 1895 experienced greater urbanization since? We expect that the most dramatic urbanization would have occurred on the edges of existing urban cores. We conducted supplemental modeling with available temporally-explicit nitrogen deposition and land use data to more directly link the temporal dynamics of drivers and response variables. While these data only cover a recent part of the entire historic period, the direction of effects did not qualitatively change from our central analysis (Fig. S2-4), and we found that there is substantially more variation across space than through time in these drivers (Fig. S1), i.e., the more urbanized regions were relatively more urban across time. Thus, while our analyses are correlational, it is likely that the use of static driver variables does not broadly change our interpretation.

Reduced endophyte detection probability within the oldest herbarium specimens, potentially due to degradation of fungal hyphae within seeds, is another possible explanation for increasing endophyte prevalence among historic specimens. Our supplemental analysis of liberal and conservative scores indicates that our results are robust to different levels of confidence in the identification of endophytes (Fig. S5-S6), however it does not entirely rule out the possibility of declining detection probability. The age of specimens is not a likely explanation for observed trends for the following reasons: (1) we only assessed endophyte status within seeds for which we could observe intact plant cells, (2) dry, climate-controlled storage conditions of herbarium specimens generally favor good preservation, and (3) our analyses identified both increasing and decreasing trends in endophyte prevalence. There is no clear reason to expect that fungal hyphae would degrade at a faster rate than host plant cells. Taphonomic studies of degradation within herbarium specimens using seeds of known endophyte status of known ages would be particularly useful to mitigate this potential source of bias, as has been done in studies of plant fossil records (Madella and Lancelotti 2012). In addition, we identified strongly supported associations between negative trends in endophyte prevalence and environmental covariates (Fig. 4; Fig. S11). If a specimen age bias does exist, it does not universally obscure the ability to detect negative trends, and such a bias would be unlikely as an explanation for environmentally-associated patterning in negative trends.

Plant-fungal interactions are fundamental pillars of biodiversity (Fei et al. 2022) and our results provide understanding of how these common interactions respond to widespread anthropogenic stressors. Broadly, our results provide evidence that this symbiosis supports host resilience to ongoing anthropogenic change. Land-use change, specifically, has been identified as a primary driver of disturbance to ecosystems globally (Parmesan and Yohe 2003; Montràs-Janer et al. 2024), and our findings contribute to a growing body of research that considers how and under what conditions these pillars of biodiversity may crumble. Our findings demonstrate clear evidence of potential resilience of plant-fungal symbiosis in the face of global change stressors, but also evidence that these interactions may be vulnerable to disruption by stress associated with urbanizing landscapes. Changing dynamics of these symbioses are likely to have consequences from populations up through ecosystems.

## List of Supplemental Material

Includes Supplemental Methods S1, Supplemental Table S1, and Figures S1-S11

## Acknowledgements

We thank the researchers and staff members of the herbaria who facilitated our research. We are also thankful for numerous undergraduate researchers who contributed to the seed scoring database. This work was supported by funding from the National Science foundation (NSF-DEB 2208857 to Tom Miller and an NSF Postdoctoral Fellowship 2410282 to Joshua Fowler). Two anonymous reviewers greatly improved earlier versions of this manuscript.

## Competing Interests

None declared

## Author Contributions

MNT contributed to research conception, data collection, data analysis and led manuscript drafting. TEX contributed to research conception, data collection, data analysis and manuscript revisions. JCF contributed to research conception, data collection, data analysis, and led manuscript revisions.

## Data Availability

All code underlying analyses can be found at: https://github.com/joshuacfowler/Endophyte_herbarium_urbanization. Data on herbarium endophyte scores are available through https://doi.org/10.5061/dryad.rn8pk0pn0, which was combined with publicly available databases of land cover and nitrogen deposition.

## Notes

https://datadryad.org/dataset/doi:10.5061/dryad.rn8pk0pn0

https://github.com/joshuacfowler/Endophyte_herbarium_urbanization

### Competing Interest Statement

The authors have declared no competing interest.

