## Supplemental figures for "Atmospheric nitrogen deposition and anthropogenic land use linked to changing fungal endophyte prevalence in cool-season grasses"

### Supplemental Materials

#### Supplemental Methods

##### 1. Evaluating the influence of year-specific vs period normal data

The analyses described in the main text that identify relationships between anthropogenic drivers (land cover type and nitrogen deposition rates) and endophyte prevalence take the average (Normal) value of each anthropogenic driver at each collection location as covariates along with year-specific values of precipitation. These Normal values are the average of relatively recent conditions based on the availability of land cover data (1985-2020) and nitrogen deposition data (1990-2020). Each herbarium specimen in our dataset has an associated year of collection spanning 1895-2019, and the environment actually experienced by plants during their lifetime may differ from this Normal value. Previous studies have documented the value of using year-specific covariates to model species-environment relationships (Milanesi et al. 2020), but also note that static variables can provide valuable information with careful interpretation (Stanton et al. 2012). Therefore, to evaluate how the use of Normal or year-specific covariate values may influence our central conclusions, we conducted a sensitivity analysis that compared model results across these types of data.

First, we found that year-specific values of anthropogenic drivers were tightly correlated with the Normal values. This was particularly clear for agricultural and urban land cover (Fig. S1 A; Fig S1 B) which has $R^2$ values of 0.989 and 0.992 respectively. Total nitrogen deposition was also highly correlated (Fig S2 C) with an $R^2$ value of 0.669. These correlation suggests that there was significantly more variation in these drivers across collection locations than through time in this data. Locations with higher levels of agricultural or urban land cover in the year of collection also tended to have higher impacted land cover across the time period of available data. We cannot evaluate whether these relationships would hold further back in time, however we think it is a reasonable assumption that highly urban or highly agricultural locations were likely to be consistently more impacted across the entire study period than locations with low contemporary levels of impacted land cover.

The association between year-specific and Normal covariate values informs the central analyses in two ways: (1) host plants collected in a location with high Normal values of anthropogenic drivers were likely to have experienced an impacted environment even in the period before available covariate data, and (2) it motivates analysis of temporal trends in prevalence to evaluate whether endophyte prevalence changed through time at a given location (the "Prevalence Trends" model in the main text that includes collection year as a covariate). Identifying temporal trends associated with high values of anthropogenic drivers could indicate either (a) that persistent high anthropogenic impacts have contributed to change in the endophyte prevalence that was obscured in the non-temporal analysis, or (b) that these highly impacted locations were formerly less impacted and that changes in endophyte prevalence through time coincide with increasing anthropogenic impact through time. While it is possible that impacts have declined through time, particularly for nitrogen deposition which has strongly declined since the late 1990s coinciding with emissions controls (with variation across regions and forms of nitrogen (Du et al. 2014; Li et al. 2016; Du 2016; Nopmongcol et al. 2019, Benish et al. 2022)), the general trend across the 20th century was rapid increases in nitrogen deposition between the 1950s and 2000 (Gschwandtner et al. 1986; Vitousek et al. 1997; Galloway et al. 2004; Cleland and Harpole 2010). And so, it is likely that the use of Normal values of nitrogen deposition in the temporally-explicit analysis can be interpreted as representing locations that were formerly less impacted but which experienced strong increases in nitrogen deposition across the period for which the majority of our herbarium specimens were collected.

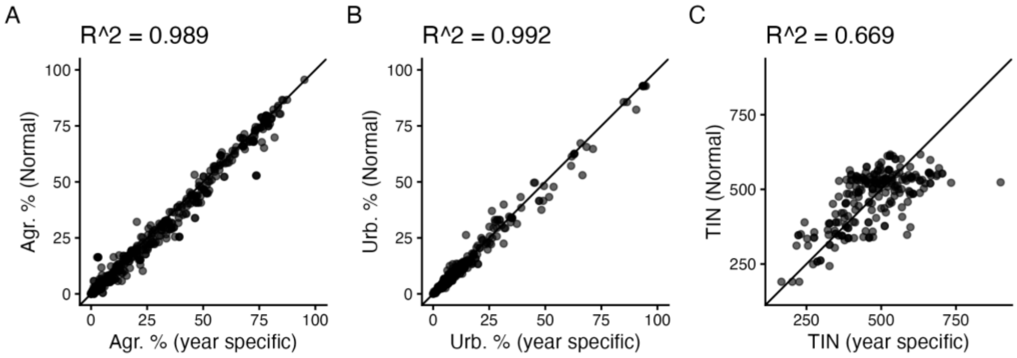


Figure S1. Correlation between year specific covariate data and Normal values taken as the average across the period of available data. (A) Percent agricultural land cover and (B) percent urban land cover data span the time period 1985-2020, and (C) Total Inorganic Nitrogen Deposition data span 1990-2020. Black points represent the values for each herbarium specimen along with the 1:1 line.

To more clearly evaluate the direct impact of year-specific or Normal data in our analyses, we compared the results of models built from the subset of data for which we had year-specific data. This necessarily reduced our data to include only 233 specimens collected between 1985 and 2019 (n = 136 *A. hyemalis*; n = 22 *A. perennans*; n = 75 *E. virginicus*). We fit models with either year-specific or Normal data, using a fixed effects structure as described in Eqn. 2 of the main text, including the effect of each anthropogenic driver along with year-specific precipitation and temperature, and with a fixed effects structure as described in Eqn. 3, including up to two-way interactions between the anthropogenic drivers, precipitation and year of collection. These models include the random effects structure described in the main text (Eqn. 1).

Comparison of model posteriors shows that broadly there is strong concurrence between parameter posterior estimates for the "static-covariate" model fit with year-specific and with Normal data (Fig. S2-S4). Notably, there was strong concordance between direction of model estimates for the main effects of anthropogenic drivers from this greatly reduced dataset and the analysis of the full dataset presented in the main text (Fig. 2). For example, the mean posterior estimate of the effect of nitrogen deposition on endophyte prevalence for *A. hyemalis* and *A. perennans* was positive. The estimated effect of nitrogen deposition for *A. hyemalis* was somewhat weaker using year-specific data than Normal data, but for *A. perennans*, the effect was stronger using year-specific data (Fig. S2). A potential explanation for this lies in the relationship between precipitation and nitrogen deposition. Annual precipitation was more strongly correlated with year-specific nitrogen deposition than average nitrogen deposition, and the model with year-specific covariates estimates a marginally stronger effect of precipitation on endophyte prevalence for *A. hyemalis*, although year-specific precipitation data was used in both models. Similar to the analyses of the main text, we estimated a positive effect of agricultural land cover on endophyte prevalence for *E. virginicus*, and a weakly negative effect of urban land cover using year-specific data (Fig. S2). Across all of these estimates, the credible intervals denote wide uncertainty overlapping zero, reflecting the significantly reduced sample size.

We repeated this comparison for the "Prevalence Trends" model, which included an effect of collection year. We again found strong overlap between parameter estimates of the model fit with year-specific as with Normal data (Fig. S3-S4). In this case, the model estimates were accompanied by much wider uncertainty and some differences in estimated direction of effects on temporal trends compared to the main analysis. For example, with this reduced dataset, the posterior estimate for the year-by-urban land cover parameter was weakly positive for *E. virginicus* (Fig. S3), while in the analysis of the main text this parameter is estimated as negative (Fig. S12). However we expect that this was largely due to the much more limited time period of sampling and smaller sample size that obscures broad temporal trends and interactions between temporal trends and anthropogenic drivers. In the "Prevalence Trends" model, estimates of main effects of anthropogenic drivers were in the same direction as the main effects of the main text analysis (Fig. S12), but the posterior estimates are largely centered around zero.

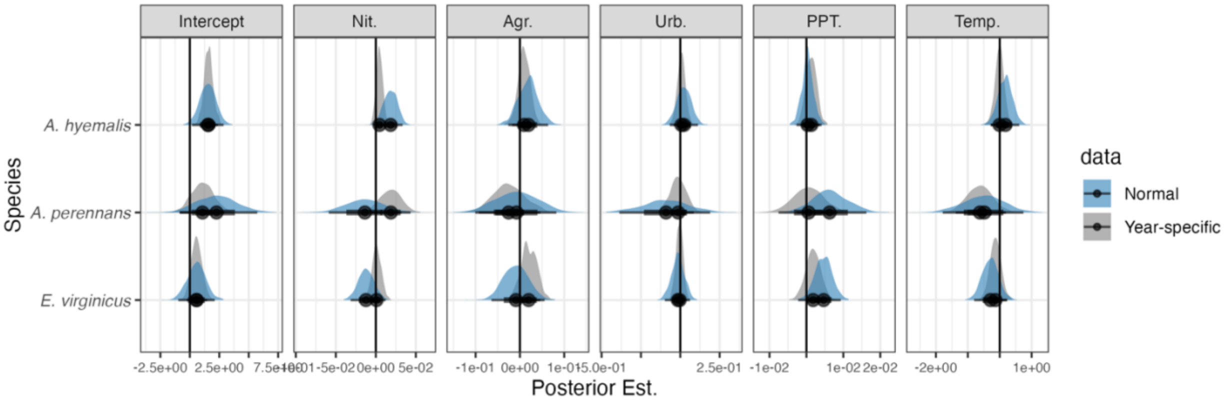


Figure S2. Parameter posterior estimates from the "Mean Prevalence" model analysis using either year-specific (grey) or period Normal data values (blue) of each anthropogenic driver on endophyte prevalence for *Agrostis hyemalis* and *Elymus virginicus*, along with the posterior mean (black circle) and 50% and 95% credible intervals.

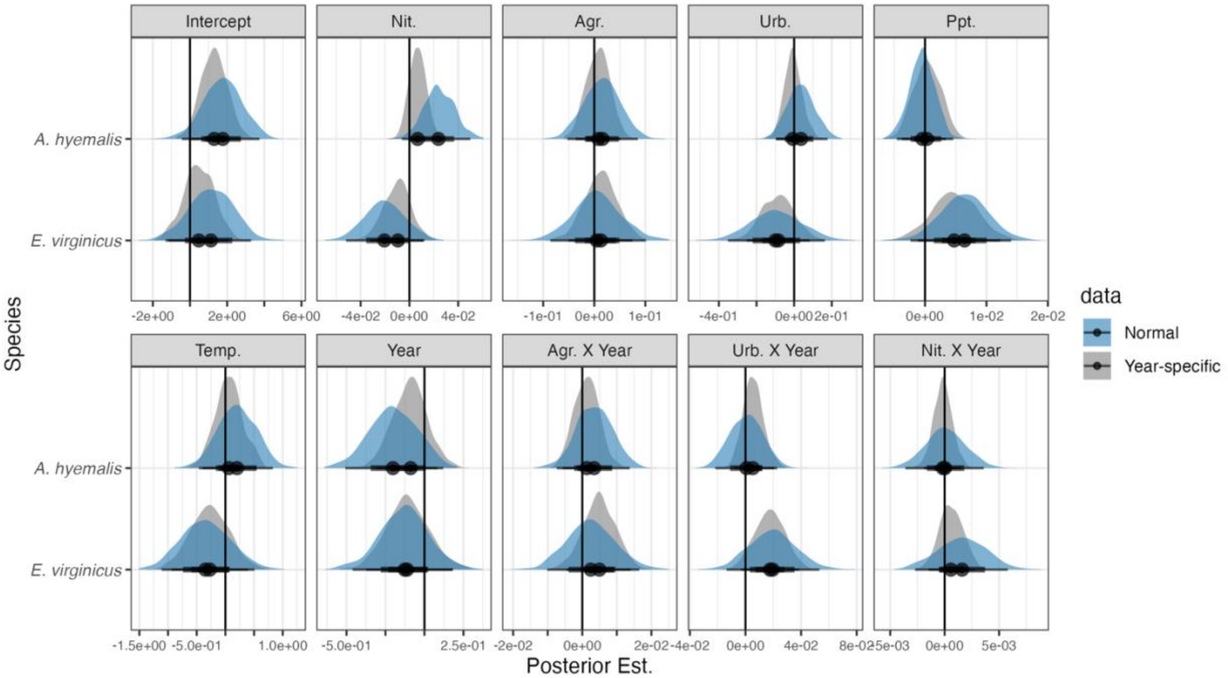

Figure S3. Parameter posterior estimates from the Prevalence Trends Model analysis using either year-specific (grey) or period Normal data values (blue) of each anthropogenic driver on endophyte prevalence for *Agrostis hyemalis* and *Elymus virginicus*, along with the posterior mean (black circle) and 50% and 95% credible intervals.

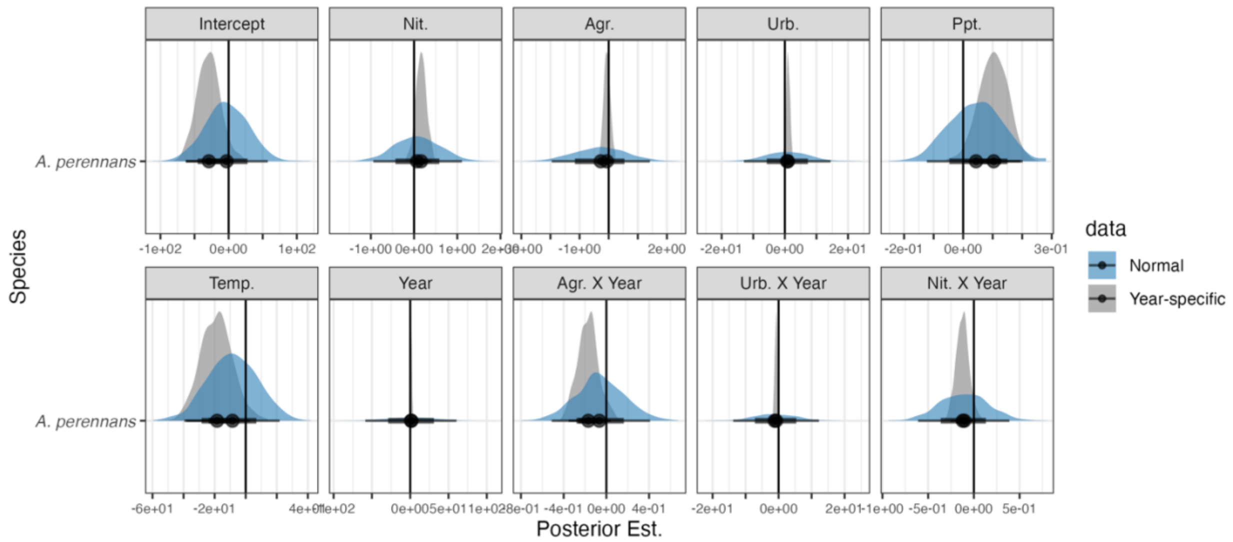


Figure S4. Parameter posterior estimates from the Prevalence Trends Model analysis using either year-specific (grey) or period Normal data values (blue) of each anthropogenic driver on endophyte prevalence for *Agrostis perennans*, along with the posterior mean (black circle) and 50% and 95% credible intervals.

#### Supplemental Figures

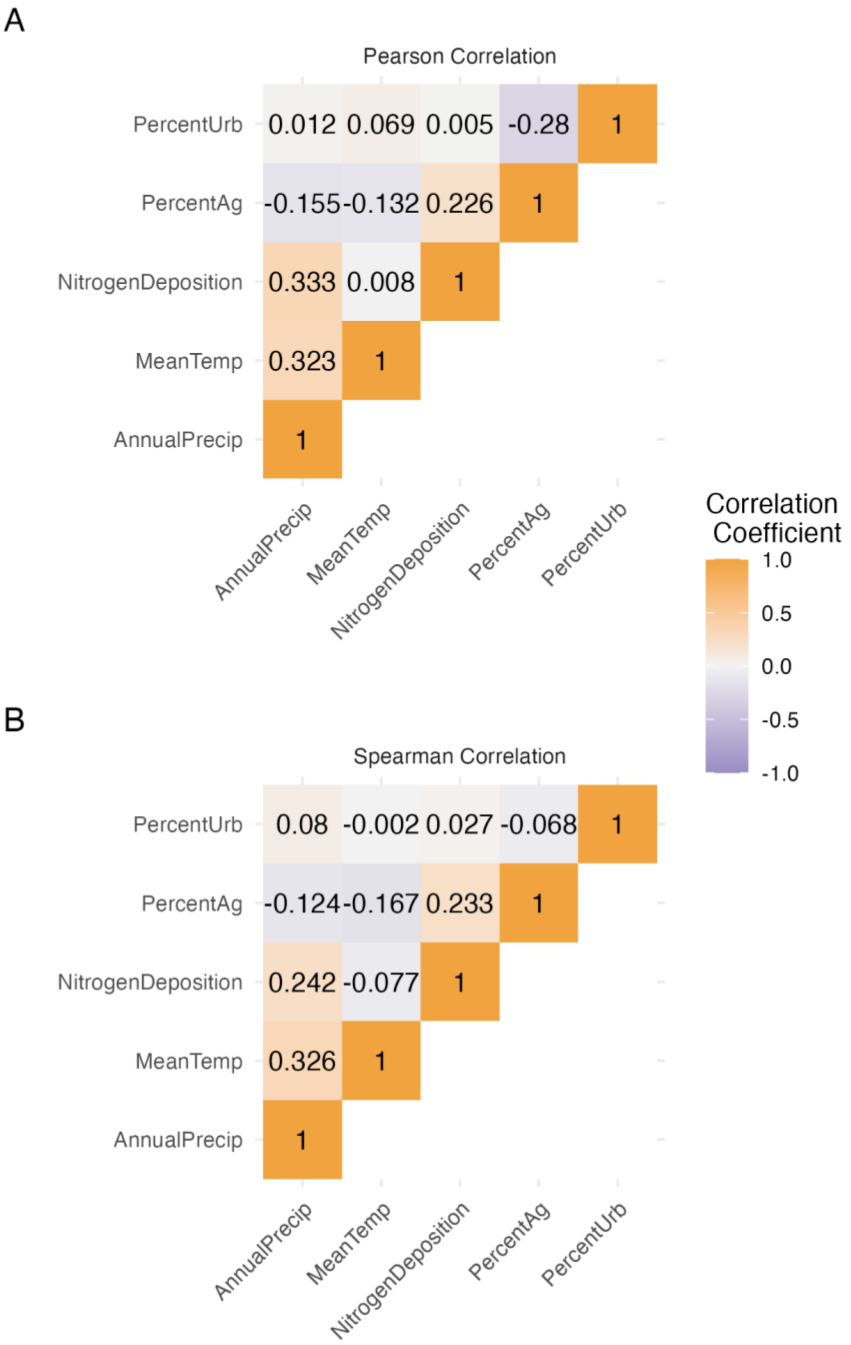


Figure S5. Correlation matrices showing (A) Pearson correlation coefficients and (B) Spearman correlation coefficients between each pairwise combination of anthropogenic global change predictor variables. Color indicates the value of the correlation coefficient.

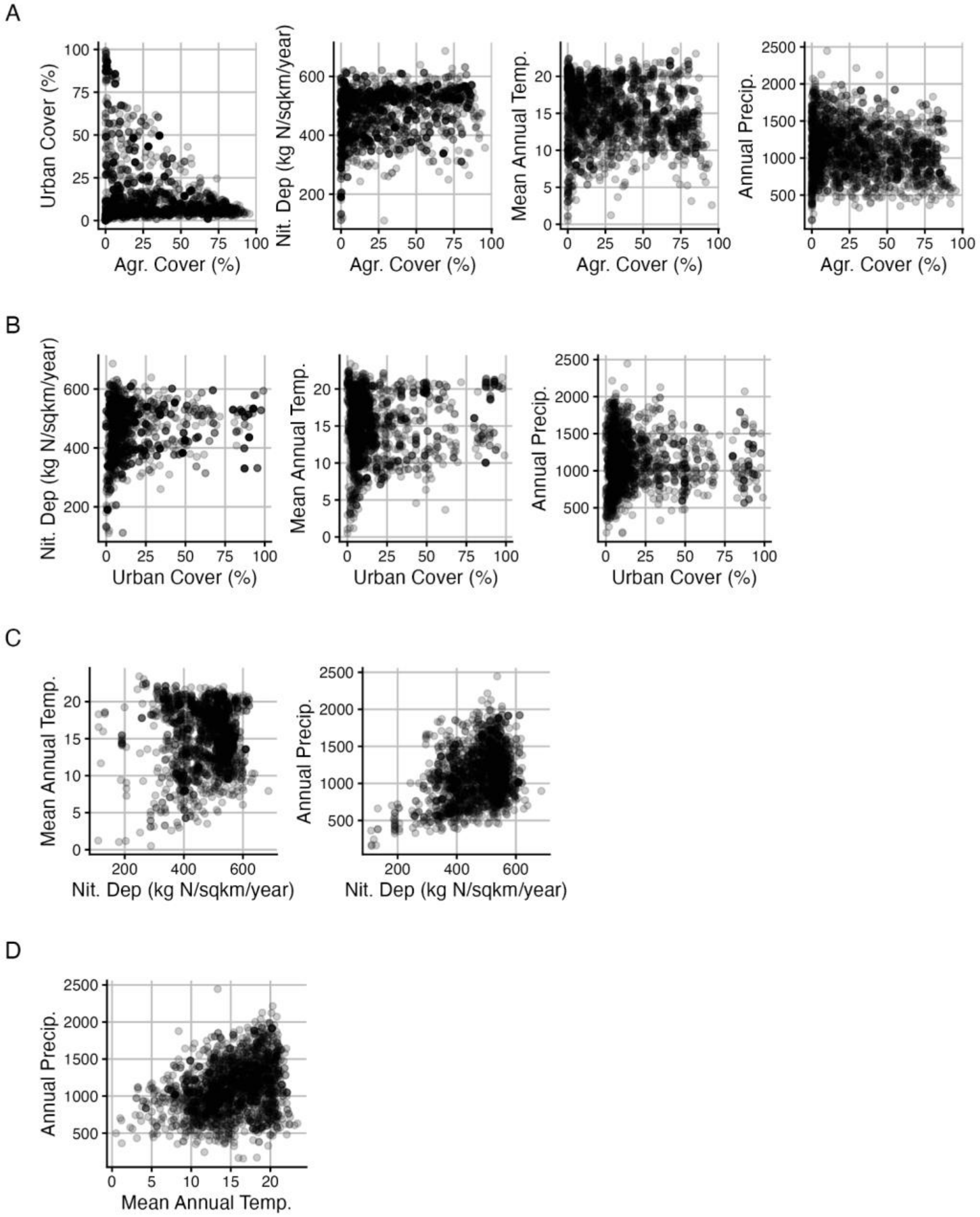


Figure S6. Pairwise visualization of global change covariate relationships. Points represent covariate values for each herbarium specimen location across (A) agricultural land cover, (B) urban land cover, (C) average annual nitrogen deposition, (D) mean annual temperature as associated with all other covariates. Land cover and nitrogen deposition values are averages of the period for which data is available (1980s-2020s), and data on climate and precipitation are annual values from the year of collection, for specimens between 1895 and 2019.

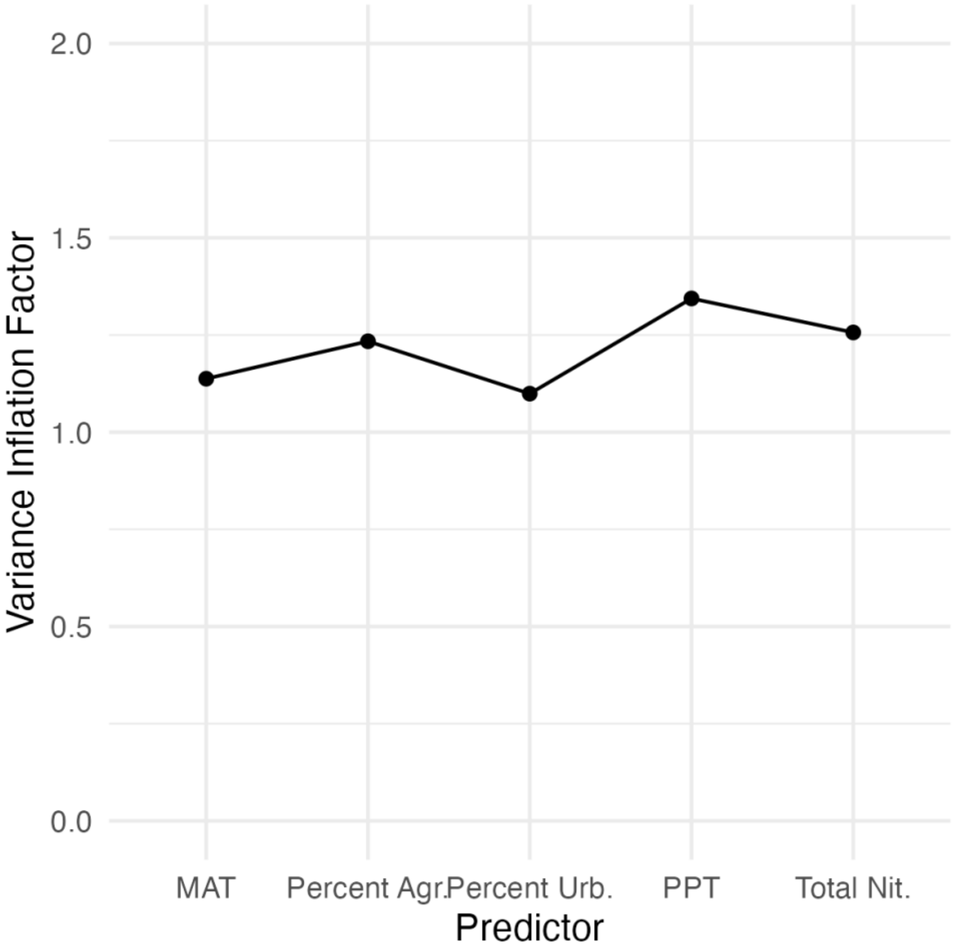


Figure S7. Variance inflation factor of each anthropogenic global change predictor variable Values close to one indicate low or no multicollinearity among these predictor variables.

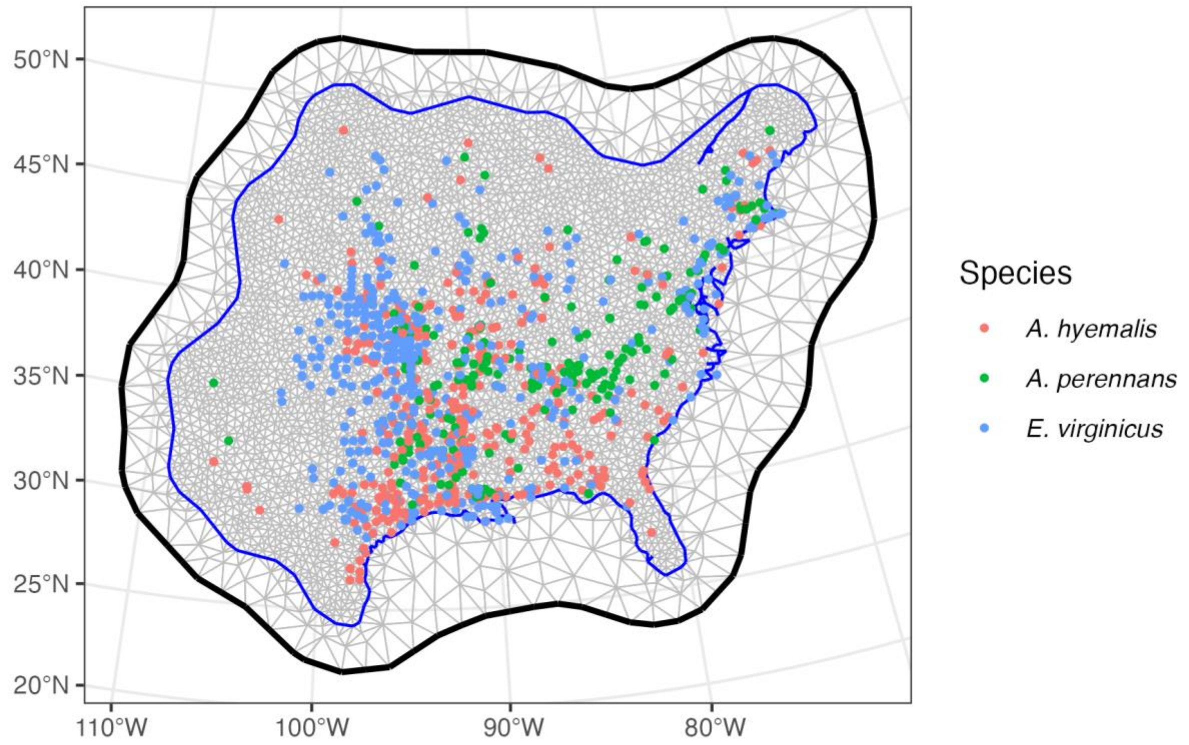


Figure S8. Delauney triangulation mesh used to estimate spatially-structured random effects accounting for potential spatial autocorrelation along with specimen collection locations for each species (red: *A. hyemalis*; green: *A. perennan*s; blue: *E. virginicus*)

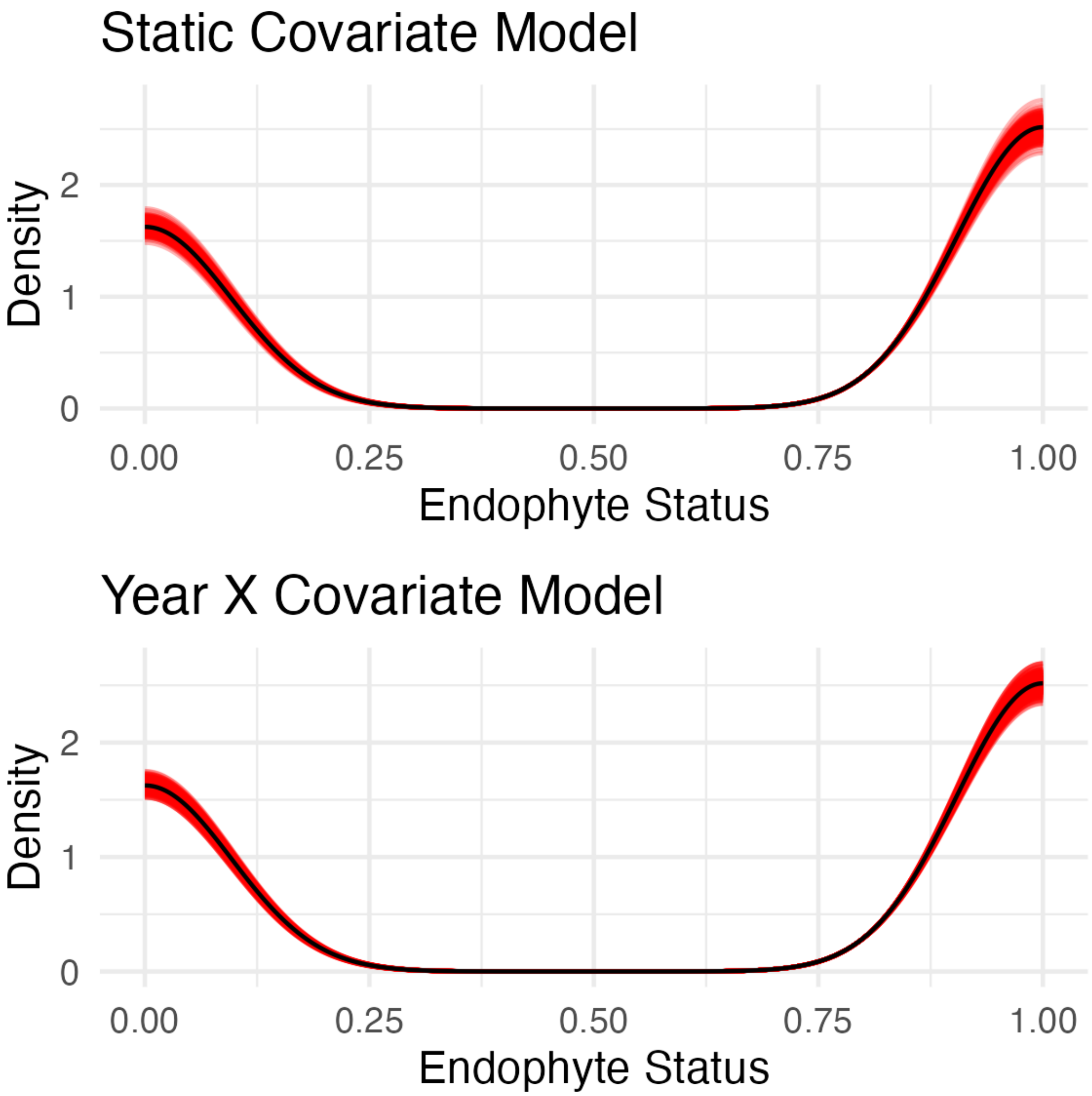


Figure S9. Posterior predictive check of the static anthropogenic driver model shows consistency between model predictions (red) and observed data (black). Lines are density curves showing predictions from 250 posterior samples. Model AUC: 0.7836.

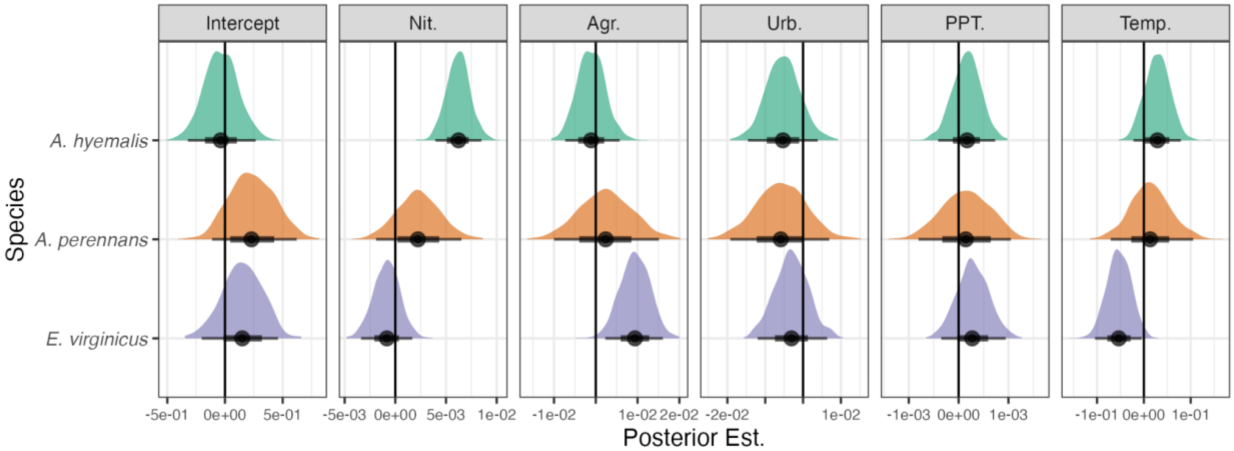


Figure S10. Parameter posterior estimates from the Mean Prevalence Model showing the effects of each anthropogenic driver on endophyte prevalence. Density curves show the sampled posterior distribution of each parameter for each species (green: *A. hyemalis*, orange: *A. perennans*, purple: *E. virginicus*) along with the posterior mean (black circle) and 50% and 95% credible intervals.

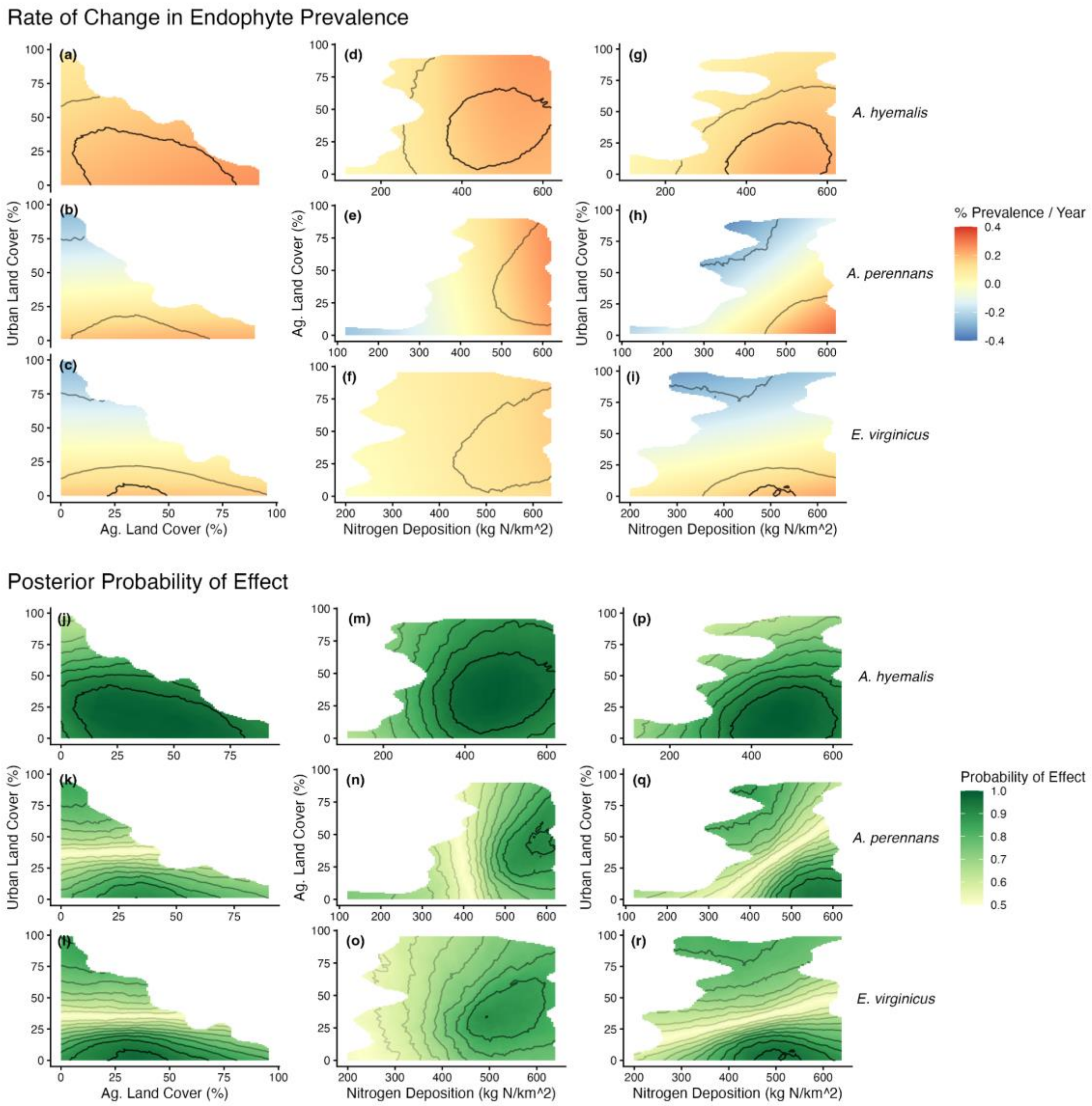


Fig. S11: Anthropogenic drivers mediate temporal trends in endophyte prevalence. Shaded regions cover the covariate space occupied by sampled herbarium specimens (e.g., there are no samples with both high urban land cover and high agricultural land cover). Shading n panels a-i depicts the mean rate of change per year in endophyte prevalence associated with anthropogenic global change drivers for *A. hyemalis* (a, d, g), *A. perennans* (b, e, h), and *E. virginicus* (c, f, i) across all levels of each anthropogenic driver. Contour lines show the region where trends have a greater than 80% (light) and 95% (dark) posterior probability of effect. Shading in panels j-r depicts the probability of effect (probability that an effect is either greater or less than 0) for *A. hyemalis* (j, m, p), *A. perennans* (k, n, q), and *E. virginicus* (l, o, r) across all levels of each anthropogenic driver. Intensity of contour lines represents increasing probability of effect from 50% to 90% posterior probability by increments of five.

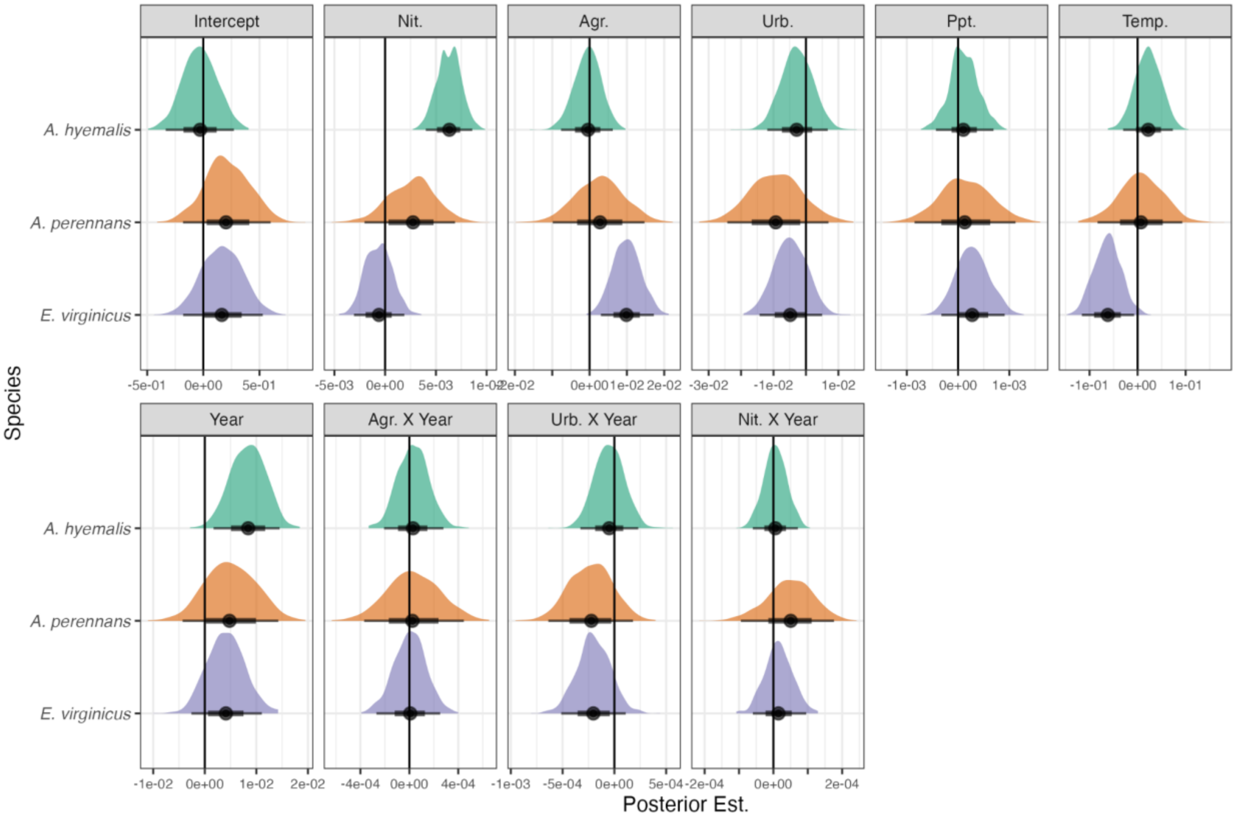


Figure S12. Parameter posterior estimates from the Prevalence Trends Model showing the effects of each anthropogenic driver on endophyte prevalence, along with interaction effects showing the effect of anthropogenic drivers in mediating temporal trends. Density curves show the sampled posterior distribution of each parameter for each species (green: *A. hyemalis*, orange: *A. perennans*, purple: *E. virginicus*) along with the posterior mean (black circle) and 50% and 95% credible intervals.

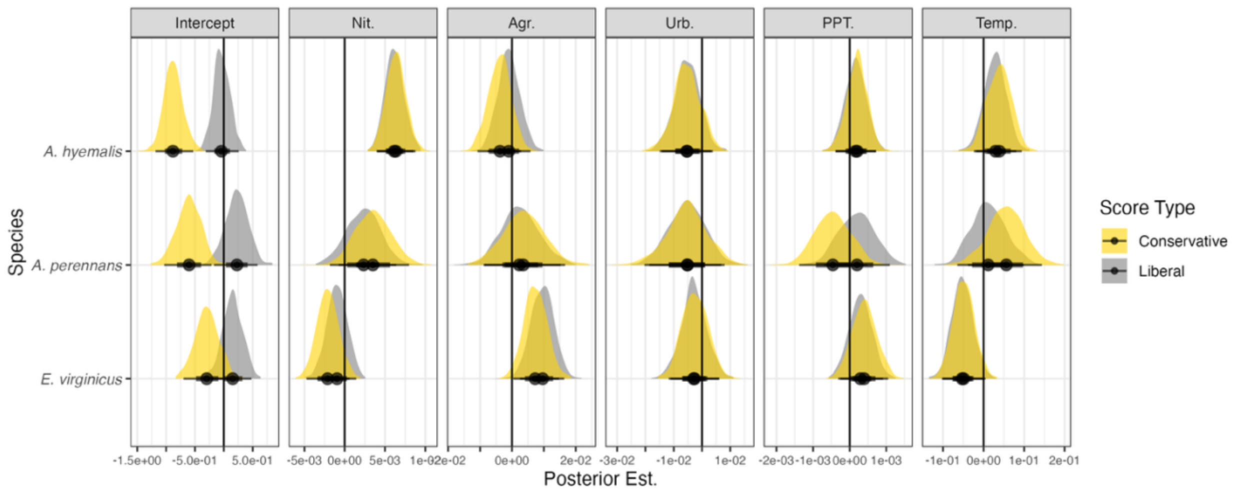


Figure S13. Parameter posterior estimates from the Mean Prevalence Model from models fit to either Conservative (yellow) or Liberal (grey) endophyte scores, representing uncertainty in the scoring process. Density curves show the sampled posterior distribution of each parameter for each species (*A. hyemalis*, *A. perennans*, *E. virginicus*) along with the posterior mean (black circle) and 50% and 95% credible intervals. The main text presents results of analysis using Liberal endophyte scores.

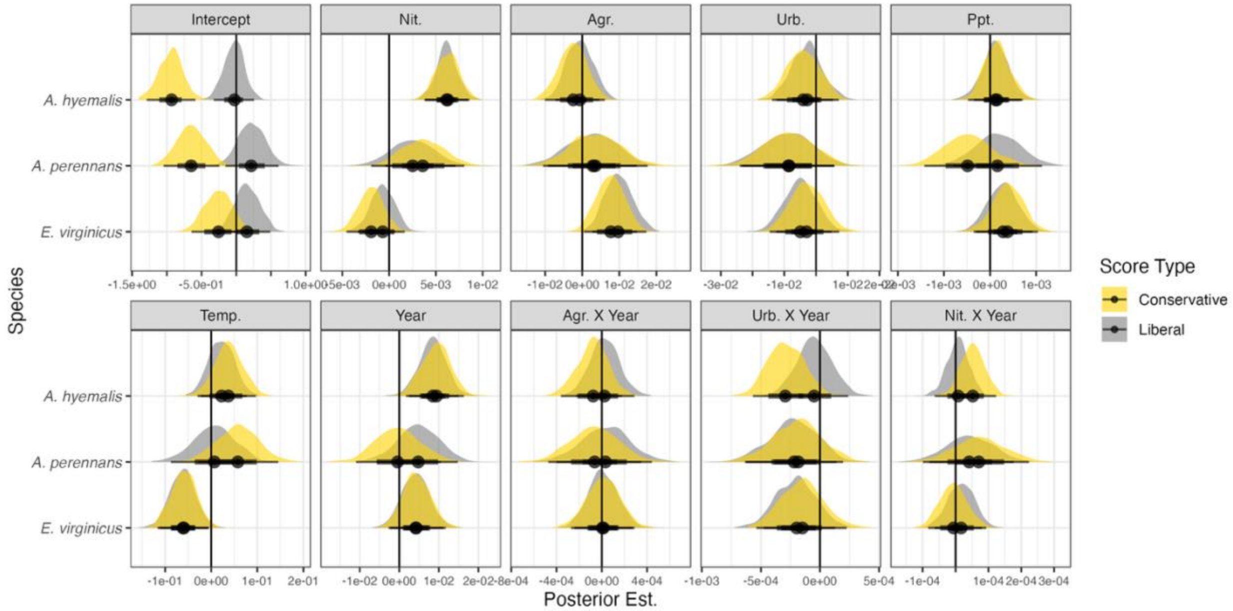


Figure S14. Parameter posterior estimates from the Prevalence Trends Model from models fit to either (a-h) Conservative (yellow) or (i-p) Liberal (grey) endophyte scores, representing uncertainty in the scoring process. Density curves show the sampled posterior distribution of each parameter for each species (*A. hyemalis*, *A. perennans*, *E. virginicus*) along with the posterior mean (black circle) and 50% and 95% credible intervals. The main text presents results of analysis using Liberal endophyte scores.

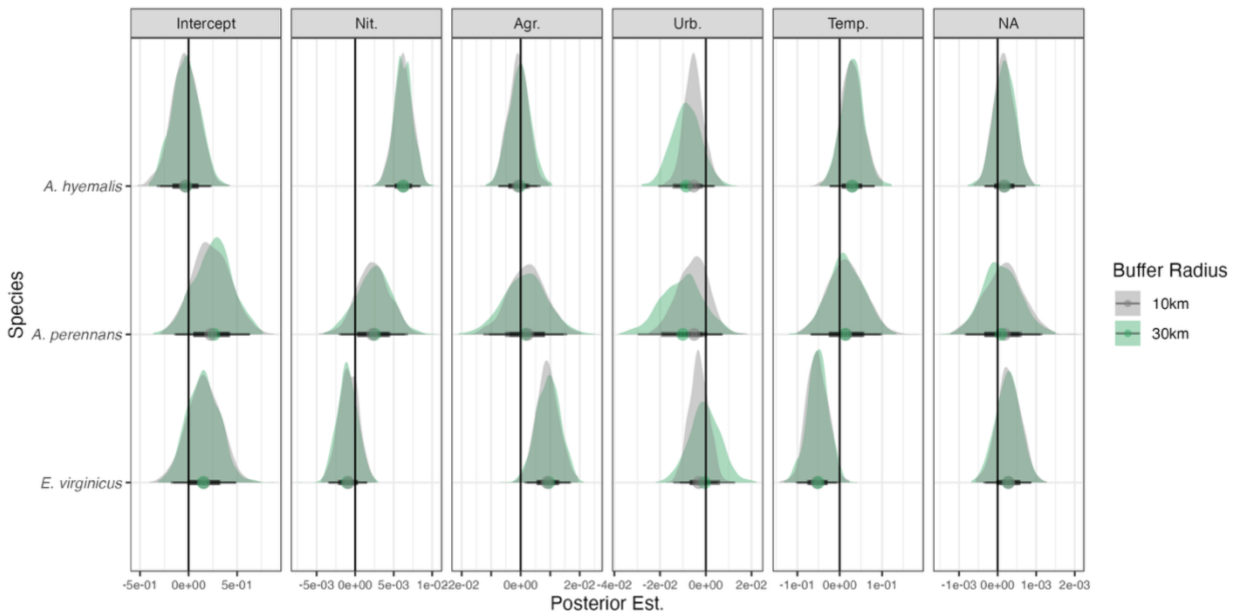


Figure S15. Parameter posterior estimates from the Mean Prevalence Model from models fit to covariate data sampled as either 30 km buffer (green) or 10 km buffer (grey). Density curves show the sampled posterior distribution of each parameter for each species (*A. hyemalis*, *A. perennans*, *E. virginicus*) along with the posterior mean (black circle) and 50% and 95% credible intervals. The main text presents results of analysis using the 10 km buffer.

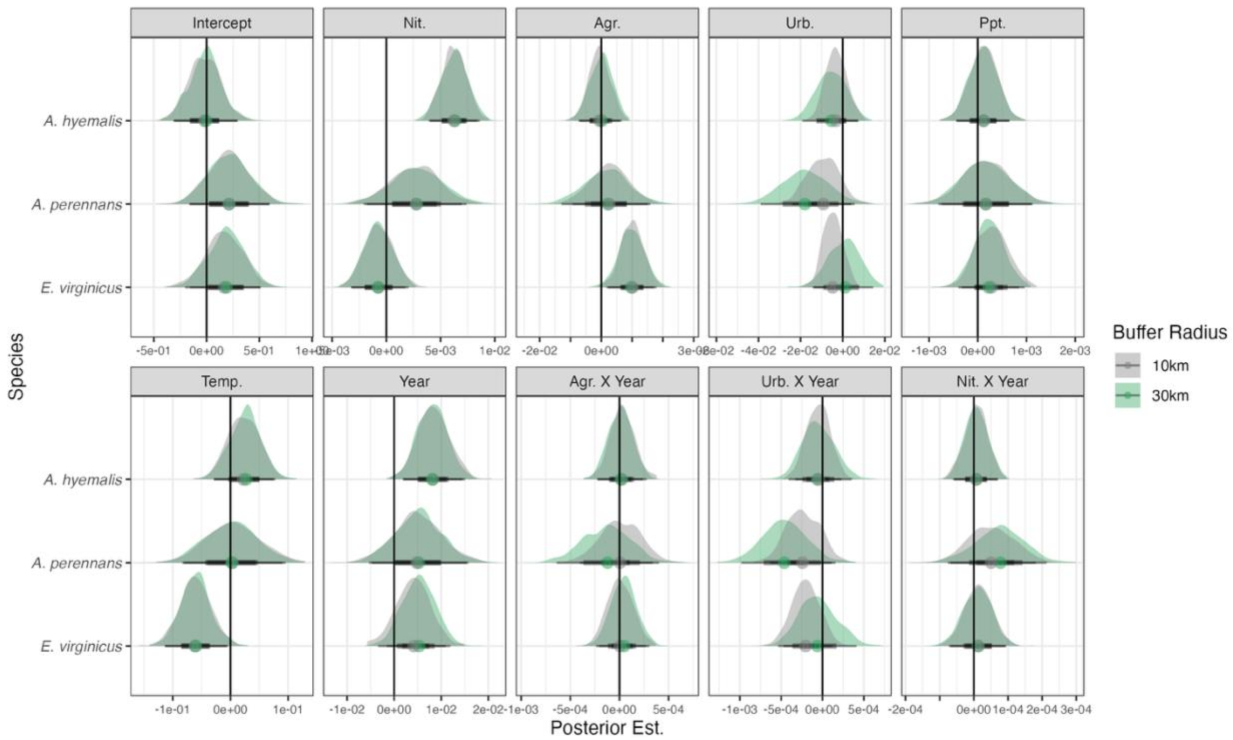


Figure S16. Parameter posterior estimates from the Prevalence Trends Model from models fit to covariate data sampled with either a 30 km buffer (green) or 10 km buffer (grey). Density curves show the sampled posterior distribution of each parameter for each species (*A. hyemalis*, *A. perennans*, *E. virginicus*) along with the posterior mean (black circle) and 50% and 95% credible intervals. The main text presents results of analysis using the 10 km buffer.

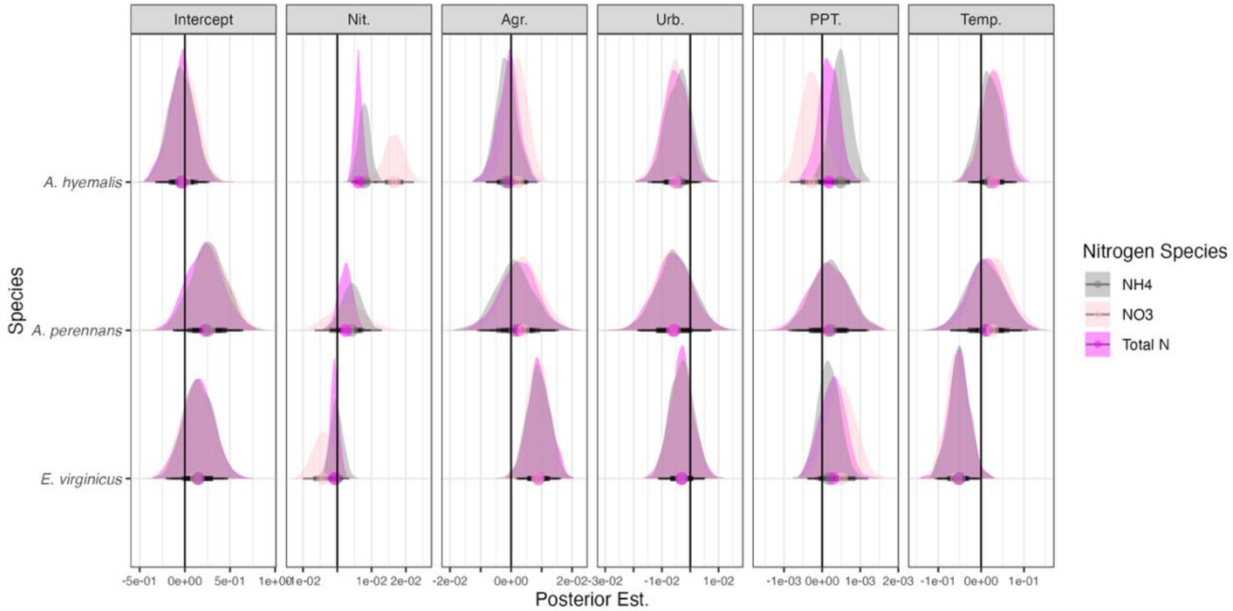


Figure S17. Parameter posterior estimates from the Mean Prevalence Model from models fit using data comprising different forms of nitrogen: NH4 (grey), NO3 (light pink), or Total inorganic nitrogen (dark pink). Density curves show the sampled posterior distribution of each parameter for each species ( *A. hyemalis*, *A. perennans*, and *E. virginicus*) along with the posterior mean (colored point) and 50% and 95% credible intervals. The main text presents results of analysis using data on total inorganic nitrogen (TIN).

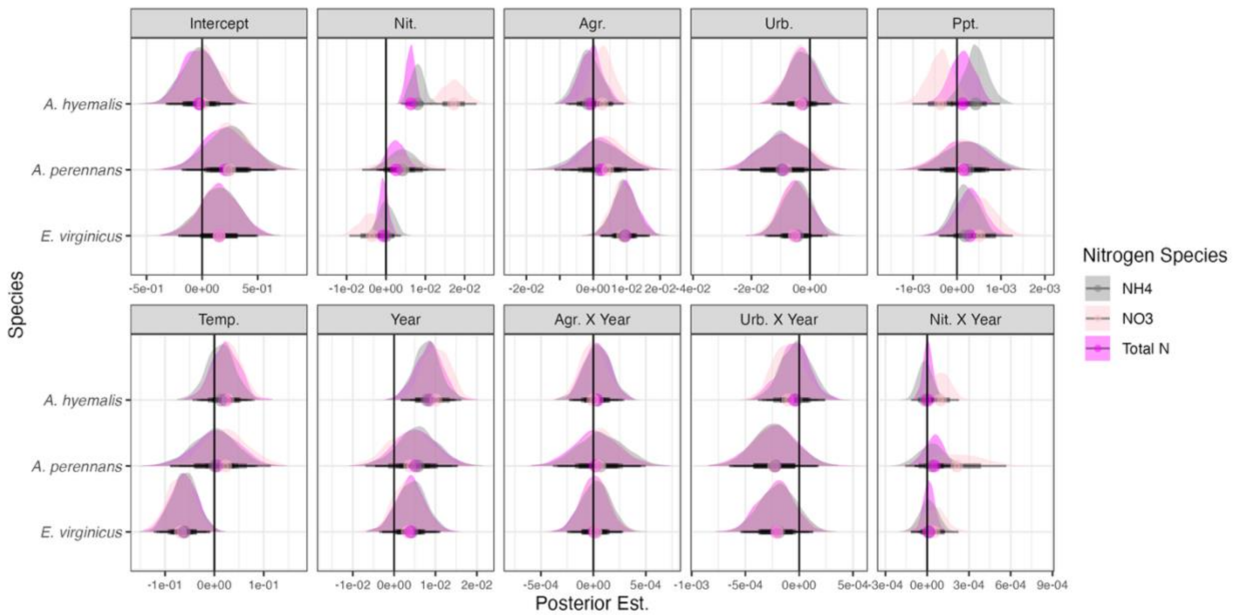


Figure S18. Parameter posterior estimates from the Prevalence Trends Model from models fit using data comprising different forms of nitrogen: NH4 (grey), NO3 (light pink), or Total inorganic nitrogen (dark pink). Density curves show the sampled posterior distribution of each parameter for each species ( *A. hyemalis*, *A. perennans*, and *E. virginicus*) along with the posterior mean (colored point) and 50% and 95% credible intervals. The main text presents results of analysis using data on total inorganic nitrogen (TIN).

#### Supplemental Tables

**Table S1.** List of herbarium collections from which seed samples were collected and the number of specimens for each species.

| **Herbarium** | **Abbreviation** | **Number of Specimens (*Elymus virginicus*)** | **Number of Specimens (*Agrostis hyemalis*)** | **Number of Specimens (*Agrostis perennans*)** |
|---|---|---|---|---|
| Botanical Research Institute of Texas | BRIT, VDB, SMU, NLU | 198 | 340 | 189 |
| Lundell Herbarium at the University of Texas at Austin | LL, TEX | 109 | 183 | 93 |
| Oklahoma State University Herbarium | OKL | 94 | 84 | 32 |
| Robert Bebb Herbarium at the University of Oklahoma | OKLA | 51 | 51 | 9 |
| Missouri Botanical Garden | MO | 120 | 211 | 144 |
| Ronald L. McGregor Herbarium at the University of Kansas | KANU | 197 | 138 | 31 |
| Shirley C. Tucker Herbarium at Louisiana State University | LSU | 61 | 72 | 38 |
| S.M. Tracy Herbarium at Texas A&M University | AM | 57 | 75 | 0 |
| Mercer Botanic Gardens | MERCA | 6 | 3 | 0 |
